# A multi-scale model of the yeast chromosome-segregation system

**DOI:** 10.1101/299487

**Authors:** Cai Tong Ng, Li Deng, Chen Chen, Hong Hwa Lim, Jian Shi, Uttam Surana, Lu Gan

## Abstract

In dividing cells, depolymerizing spindle microtubules move chromosomes by pulling at their kinetochores. While kinetochore subcomplexes have been studied extensively *in vitro*, little is known about their *in vivo* structure and interactions with microtubules or their response to spindle damage. Here we combine electron cryotomography of serial cryosections with genetic and pharmacological perturbation to study the yeast chromosome-segregation machinery at molecular resolution *in vivo*. Each kinetochore microtubule has one (rarely, two) Dam1C/DASH outer-kinetochore assemblies.

Dam1C/DASH only contacts the flat surface of the microtubule and does so with its flexible “bridges”. In metaphase, 40% of the Dam1C/DASH assemblies are complete rings; the rest are partial rings. Ring completeness and binding position along the microtubule are sensitive to kinetochore attachment and tension, respectively. Our study supports a model in which each kinetochore must undergo cycles of conformational change to couple microtubule depolymerization to chromosome movement.

## INTRODUCTION

The spindle apparatus, a microtubule-based machine, partitions chromosomes equally between mother and daughter cells during mitosis. In yeast, the microtubules (MTs) in both the nucleus and cytoplasm are anchored by their closed “minus” ends to the nuclear-envelope-embedded microtubule-organizing centers, termed spindle pole bodies. The MT “plus” end (the tips of the 13 protofilaments) can have either a flared or ‘ram’s horn’ configuration (Winey et al., 1995). Kinetochore MTs (kMTs) attach to chromosomes while the long pole-to-pole MTs render the spindle its characteristic shape. To prevent chromosome missegregation, cells employ the spindle assembly checkpoint (SAC) to delay anaphase onset until two conditions are met: first, each sister chromosome must attach to kMTs emanating from one of the spindle pole bodies (bi-orientation or amphitelic attachment) (Musacchio and Salmon, 2007). Second, the spindle must generate tension via opposition between the kMT-induced poleward pulling forces and the cohesion between sister chromatids mediated by cohesin complexes (Michaelis et al., 1997). Damaged spindles and erroneous kMT attachments resulting in either unoccupied kinetochores or a loss of tension in the spindle apparatus leads to the activation of the SAC. The activated SAC imposes a transient cell-cycle arrest in prometaphase, allowing cells to restore kinetochore-microtubule attachments before progressing to anaphase (Tanaka, 2010).

The kinetochore is a multi-functional protein complex that mediates the chromosome-kMT attachment and couples kMT depolymerization to poleward movement of the chromosome. Furthermore, the kinetochore is central to the SAC because it can assess the quality of chromosome-kMT attachment. Kinetochores are so complex that its subassemblies -- classified as centromere-proximal “inner-kinetochore” or kMT-associated “outer-kinetochore” complexes based on traditional EM studies -- are often studied as reconstituted complexes (Musacchio and Desai, 2017). High-precision fluorescence imaging *in vivo* has revealed the composition and the average positions of many of these subassemblies (Joglekar and Kukreja, 2017). In yeast, the best-understood one is the outer-kinetochore Dam1C/DASH complex (Cheeseman et al., 2001; Hofmann et al., 1998; Janke et al., 2002; Jones et al., 1999; Li et al., 2002). Ten different polypeptides assemble as a Dam1C/DASH heterodecamer (Miranda et al., 2005). Dam1C/DASH heterodecamers can further oligomerize as rings around MTs (Miranda et al., 2005; Westermann et al., 2005). Owing to their circular shape and ability to form stable load-bearing attachments on MTs *in vitro* (Asbury et al., 2006; Franck et al., 2007; Westermann et al., 2006), Dam1C/DASH rings are thought to anchor the chromosome onto kMTs and couple kMT depolymerization to chromosomal poleward movement by interacting with the protofilaments’ curved tips (Efremov et al., 2007).

Knowledge of kinetochore structure at the molecular level *in vivo* would shed light on fundamental questions that cannot be addressed by reconstitution. These questions include how the kinetochores couple to the kMTs; how the kinetochore subunits are oligomerized; how kinetochores are distributed in 3-D within the spindle; and how both the kinetochore and spindle respond to perturbation. These structural details remain largely unknown *in vivo* because kinetochores are sensitive to conventional electron-microscopy sample-preparation methods (McEwen et al., 1998; McIntosh, 2005).

Structural insights into large complexes like kinetochores and spindles *in vivo* require electron cryotomography (cryo-ET), which can reveal the 3-D architecture of giant cellular machines and their subcomponents in a life-like state (Gan et al., 2011).

We used cryo-ET of both serial and single frozen-hydrated sections (cryosections) to test decades-old structural models of the yeast chromosome-segregation system *in vivo*. We have examined the structure of yeast outer-kinetochore Dam1C/DASH oligomers and their interactions with kMT walls in metaphase cells both with and without tension, in cells treated with a spindle poison, and in comparison to Dam1C/DASH-MT complexes *in vitro*. We found that Dam1C/DASH can oligomerize into two types of rings, both of which can stably associate with kMTs. Finally, our study reconciles different views concerning the mechanism of outer-kinetochore function in a new model of MT-powered chromosome movement.

## RESULTS

### Dam1C/DASH forms both complete and partial rings *in vitro*

To understand how individual Dam1C/DASH rings interact with MTs, we performed cryo-ET of plunge-frozen Dam1C/DASH assembled around MTs *in vitro* (Fig. S1) (Miranda et al., 2005). We observed both complete and partial rings (Fig. 1A-B).

**Figure 1.**
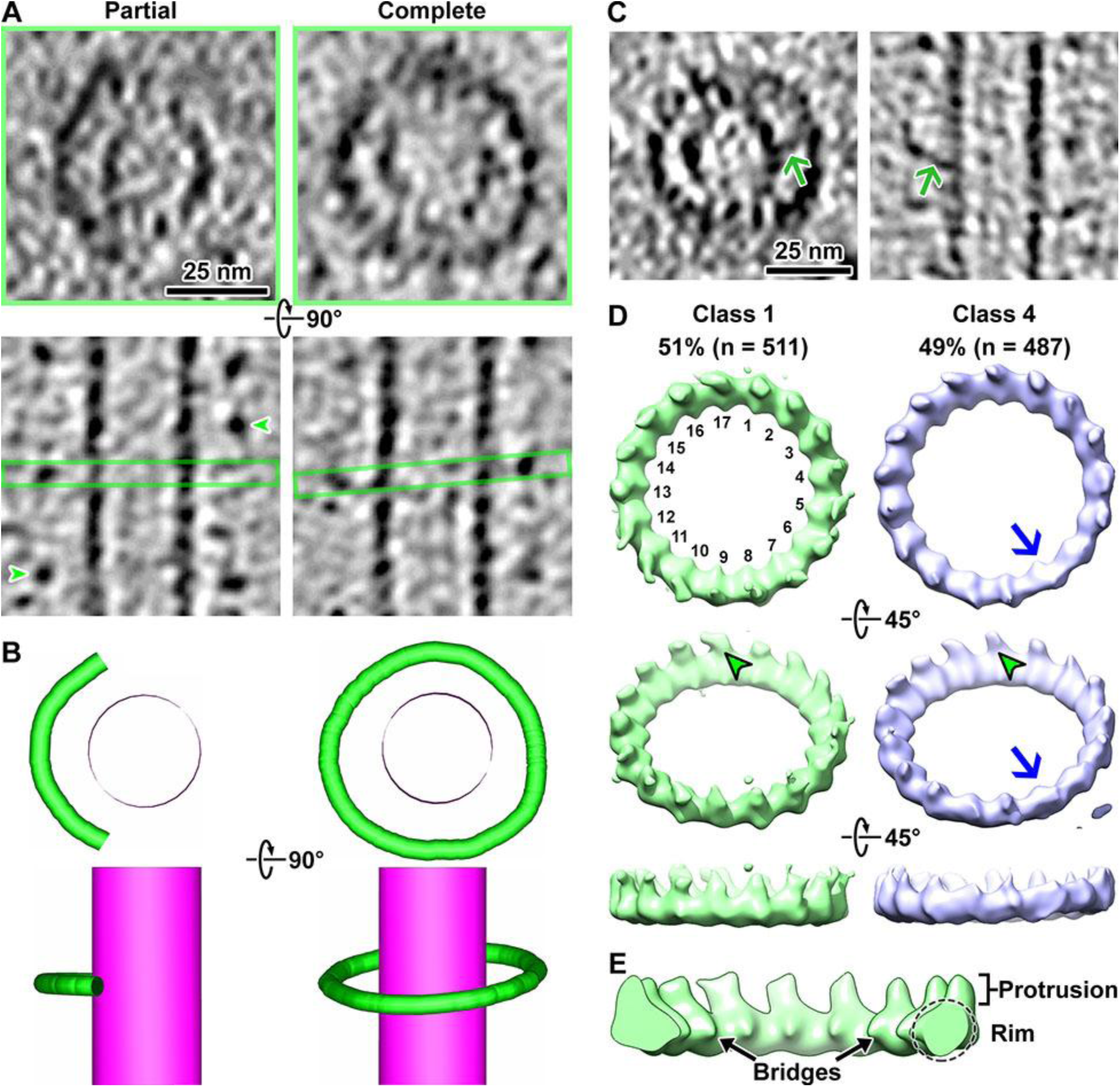
Dam1C/DASH oligomerizes into partial and complete rings *in vitro*. (A) Cryotomographic slices (4.6 nm) showing front views of partial (left) and complete (right) Dam1C/DASH rings assembled around MTs. The lower row shows the same rings but rotated 90° around the horizontal axis. Green arrowheads: densities of adjacent Dam1C/DASH oligomers; green rectangles: approximate planes of the partial or complete ring taken in the upper panels. (B) Three-dimensional models of Dam1C/DASH and MT complexes corresponding to upper and lower rows in panel A. (C) Two examples of Dam1C/DASH rings with bridges (green arrows), in the front (left) and side (right) views. (D) Asymmetric 3-D class averages of Dam1C/DASH rings around MTs. Repeat subunits are numbered for class 1. Classes 2 and 3 (not shown) are very similar to class 4 and were included in the 49%. The upper image is the front view. The middle and lower panels are sequentially rotated 45° around the horizontal axis. Green arrowheads: protrusions. Blue arrow: position in class average 4 that deviates from 17-fold symmetry. All density maps were masked to exclude the MT and contoured at 0.1 σ above the mean. (E) Enlarged, cutaway view of a 17-fold symmetrized Dam1C/DASH ring, with landmark motifs labeled. Note that the bridges appear shorter than in the individual subtomograms because their structures are extremely heterogeneous.

Consistent with previous studies (Miranda et al., 2005; Westermann et al., 2005), our cryotomograms showed that most Dam1C/DASH rings are slightly tilted relative to the MT’s axis. Furthermore, most of these complete and partial rings have flexible structures that connect the ring’s rim to the MT walls (Fig. 1C). These connections are called “bridges” (Miranda et al., 2007; Wang et al., 2007; Westermann et al., 2005) and are thought to be composed of parts of the Dam1p and Duo1p proteins (Legal et al., 2016; Zelter et al., 2015).

Rotational power-spectra analyses (Murphy et al., 2006) of individual rings showed that most of the complete Dam1C/DASH rings had 17-fold symmetry *in vitro* (Fig. S2A-G). This conclusion was further supported by asymmetric 3-D class averages, which also have 17-fold symmetry (Fig. 1D and S3). Unlike in previous studies (Ramey et al., 2011; Wang et al., 2007; Westermann et al., 2006), we did not observe any 16-fold-symmetric rings *in vitro*; the reason for this difference is not clear. Nevertheless, our Dam1C/DASH structure shares similar motifs with the previous ring structure (Ramey et al., 2011), such as the inward-pointing stump-like densities that correspond to a portion of the bridge (Fig. 1E) and densities extending from the ring’s rim, parallel with the MT surface. We call these latter motifs “protrusions”, following the nomenclature of a recent structure of reconstituted Dam1C/DASH rings (Steve Harrison, personal communication). For brevity, herein we use the terms bridge, rim, and protrusion when referring to these prominent Dam1C/DASH structural motifs (Fig. 1E).

### Strategy to study yeast kinetochore structure *in vivo*

The *in vivo* structure of Dam1C/DASH is unknown. Previous tomography studies of high-pressure-frozen, freeze-substituted cells revealed weak densities at kMT plus ends that might be partial or complete rings (McIntosh et al., 2013). To eliminate fixation, dehydration and staining as sources of structural distortion, we prepared all our cells by high-pressure freezing, followed by thinning to ~ 100 - 150 nm by cryomicrotomy. As a positive control, we assembled Dam1C/DASH rings around MTs *in vitro* and subjected these samples to the same high-pressure freezing and cryomicrotomy done for cells.

The contrast of cryotomograms from such samples is extremely low due to the high concentration of the dextran cryoprotectant (Chen et al., 2016). Both partial and complete Dam1C/DASH rings were nevertheless visible in the resultant cryotomograms (Fig. S4). Therefore, cryo-ET of cryosections can reveal both partial and complete Dam1C/DASH rings around kMTs if they exist *in vivo*.

We prepared mitotic yeast cells with either attached kinetochores under tension, detached kinetochores without tension, or attached kinetochores without tension (Fig. 2A). Knockdown of Cdc20 function causes yeast cells to arrest in metaphase with kinetochores attached to the spindle and under tension (Lau and Murray, 2012; O’Toole et al., 1997). To visualize kinetochores in metaphase, we arrested cells by depleting Cdc20 in a GAL-Cdc20 strain (Fig. 2B). Because kinetochores take up a tiny fraction of the cell’s volume, a single cryosection taken randomly through a cell is unlikely to contain a kinetochore. To overcome this challenge, we devised a parallel-bar-grid-based serial-cryo-ET workflow that made possible the reconstruction of larger portions of spindles (Fig. 2C and D, 3A).

**Figure 2.**
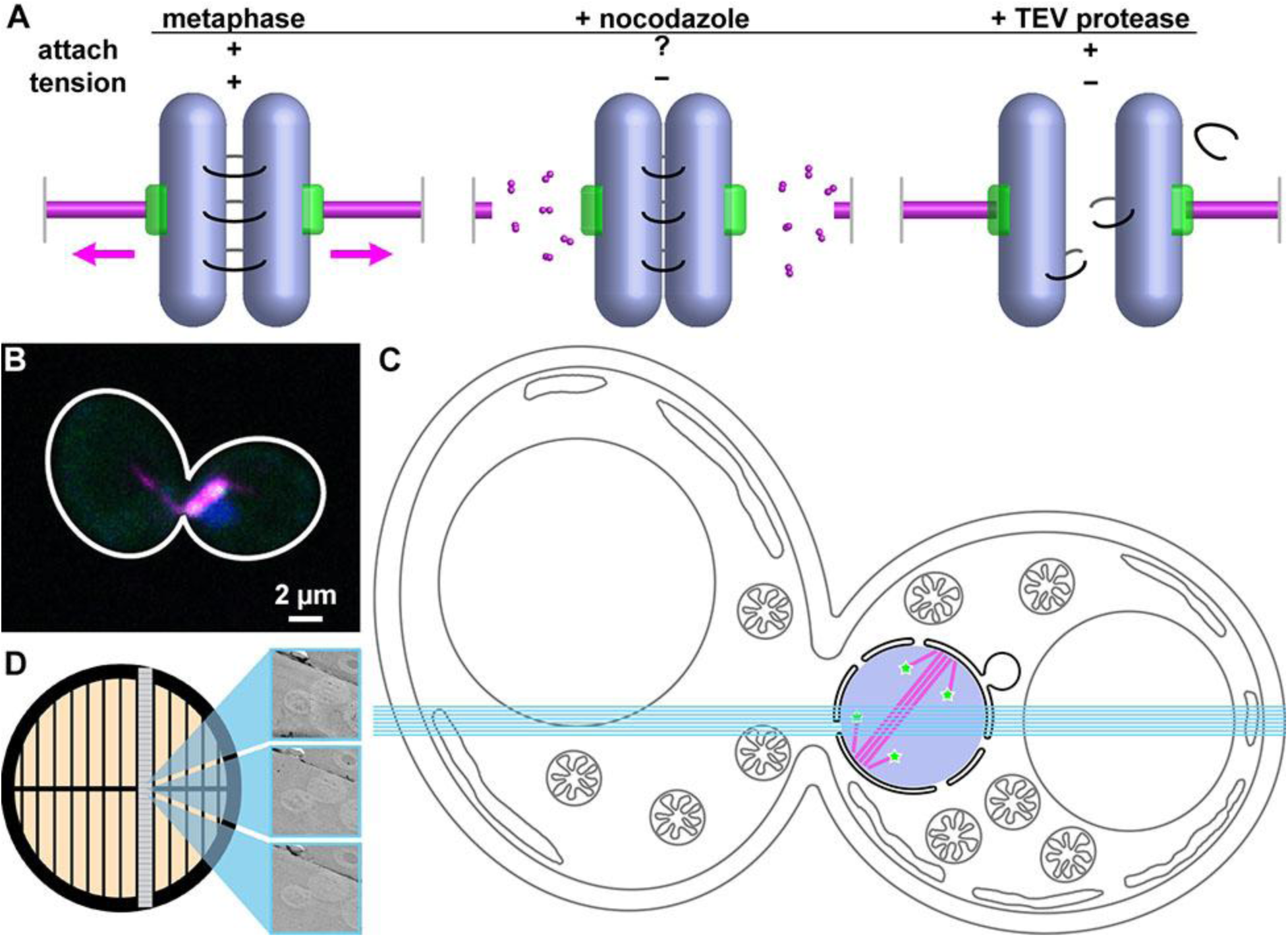
Strategy to find kinetochores *in vivo*. (A) Schematic of kinetochore states studied in this paper, not to scale. Sister chromosomes (pale-blue rods) are held under tension (magenta arrows) in metaphase when kMT (magenta tubes) pulling forces are transmitted by cohesin (curved lines). Gray vertical bars: spindle-pole bodies. Tension at the kinetochores (green) can be eliminated either by the disruption of the kMTs with nocodazole or the conditional cleavage of mutant cohesin with TEV protease. This color scheme is used throughout the paper. (B) Immunofluorescence image of a Cdc20-depleted cell, with Dam1p-GFP in green and Tubulin in magenta. Owing to the merged channels, Dam1C/DASH appears white. (C) Cartoon of a mitotic yeast cell, with organelles drawn to approximate scale. The nucleus (pale blue circle), spindle (magenta lines), and kinetochores (green stars, not to scale) are colored. Cyan lines illustrate at scale a series of seven ~ 100-nm cryosections. (D) Serial cryotomography strategy. Cryo-EM images of sequential cryosections of the same cell mounted on a parallel-bar grid are shown enlarged ~ 100-fold on the right.

### Dam1C/DASH forms both complete and partial rings around kMTs *in vivo*

We reconstructed portions of 23 metaphase spindles, each with at least 3 serial cryotomograms (most-complete example in Fig. S5). We identified kMTs based on their short length, location, and orientation relative to the nuclear envelope (Fig. 3B). Both complete and partial ring structures encircled the kMT plus ends (Fig. 3C-F). We herein assign these complete and partial rings as Dam1C/DASH because their shape, diameter (47 ± 5 nm, mean and standard deviation, n = 12), localization at the kMT plus-ends, bridge densities (see below), and their absence from cytoplasmic MTs (see below) are all consistent with that expected of Dam1C/DASH from *in vivo* and *in vitro* studies. Our most complete serial-cryo-ET reconstruction (Figs. 3B and S5) contained half a spindle with 13 Dam1C/DASH rings (Fig. 3B). Budding yeast cells have 16 kinetochores per half spindle (one kinetochore per sister chromosome), so we probably missed 3 Dam1C/DASH rings due to the ambiguity of cryo-ET densities near the cryosection surfaces (14 surfaces in 7 cryosections). The reconstruction is complete enough that we estimate that all kinetochores would fit into a rectangular volume less than 0.5 µm on a side (Fig. 3C).

**Figure 3.**
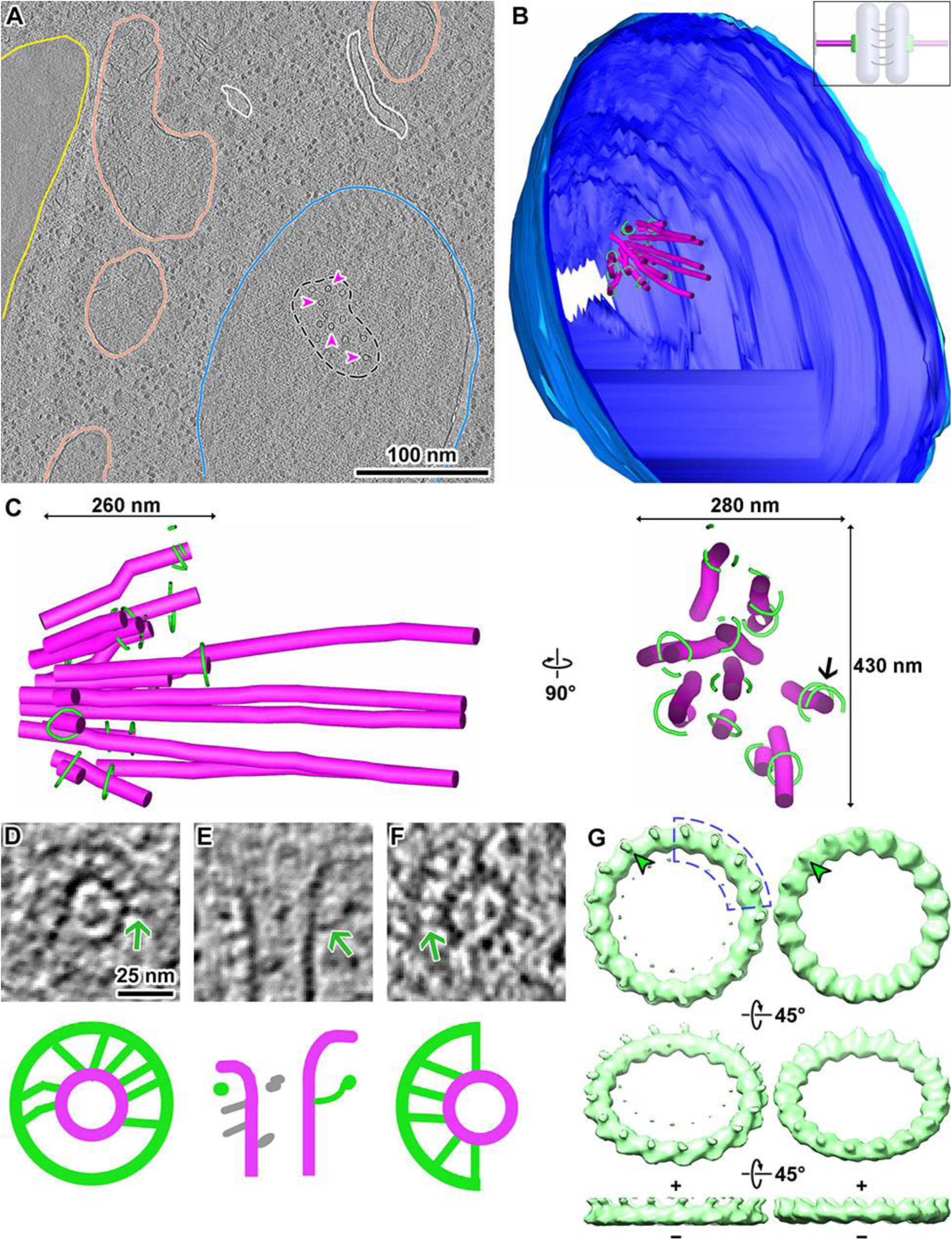
Architecture of metaphase spindles and outer kinetochores. (A) Cryotomographic slice (18 nm) from a Cdc20-depleted cell. Major features are annotated: cell membrane (yellow), mitochondria (salmon), endoplasmic reticulum (white), nucleus (blue). The black dashes outline the spindle. A few spindle MTs are indicated with magenta arrowheads. (B) Three-dimensional model of a half spindle, spanning 7 sequential sections. Dark and light blue: inner and outer nuclear membranes. Magenta tubes: spindle MTs. Green rings: Dam1C/DASH. Inset: schematic showing the structures that are modeled (saturated shading) and those that are not (washed-out shading). (C) Left: Enlargement of the spindle modeled in panel B and rotated to a view perpendicular to the spindle’s axis. Right: Transverse view of the same spindle; for clarity, polar MTs are omitted. Because the short axis crosses multiple cryosection interfaces, we are uncertain how long this spindle was in the unsectioned cell. This particular spindle also has an oval cross section due to microtomy compression along the X axis of the right panel. Black arrow: one example of a kMT with two partial rings. (D - F) Cryotomographic slices (6 nm) of Dam1C/DASH rings around kMTs. Green arrows point to bridges. The lower panels show schematics of the Dam1C/DASH (green), kMT (magenta), and kMT-associated protein (gray) densities. Panels D and F show front views of a complete and partial ring, respectively. Panel E shows a side view of a complete ring. (G) Rotationally averaged density maps of two individual complete Dam1C/DASH rings *in vivo*, masked to exclude the kMT, contoured at 1σ above the mean. Top: front view. The middle and lower rows are sequentially rotated 45° around the horizontal axis. Green arrowheads: protrusions. The plus and minus signs indicate the polarity of the encircled kMT. If four or more decamers (outlined by blue dashes) were absent, there would be a gap > 25 nm.

### Dam1C/DASH bridges contact the flat or gently curved surfaces of kMTs

Most kMTs had a single Dam1C/DASH ring (complete or partial, n = 82) (Fig. 3B and C). Only 4 kMTs had two partial Dam1C/DASH rings each (one example shown in Fig. 3C). All rings were tilted relative to the kMT axis and the majority of them were positioned within 50 nm of the plus end. We did not observe any contacts between Dam1C/DASH and the protofilaments’ curved tips. Instead, the only Dam1C/DASH-kMT interactions observed were between the kMT walls and Dam1C/DASH’s bridges (Fig. 3D-F; additional examples in Fig S6). Furthermore, we did not observe any contact between Dam1C/DASH and the back of the protofilaments’ curve tips *in vitro* (Fig. S7). The Dam1C/DASH bridges are conformationally heterogeneous, even within the same ring, and could either be coplanar with the rim or curved out of plane (Fig. 3D and E). If each Dam1C/DASH heterodecamer contributes a single bridge, then there would be up to 17 bridges per ring. In both our *in vivo* and *in vitro* datasets, we observed up to 8 bridges per ring, meaning that most of the bridges were in an as-yet-unknown conformation. This conformational flexibility explains how the bridge appears as a continuous density from the Dam1C/DASH rim to the MT surface in cryotomograms but as a stump-like density in multi-ring averages.

Two complete Dam1C/DASH rings *in vivo* had sufficient contrast to reveal that they had 17-fold symmetry (one analysis shown in Fig. S2H). To better understand how Dam1C/DASH is organized *in vivo*, we symmetrized these two rings, yielding density maps with higher signal-to-noise-ratio (Fig. 3G). These symmetrized rings have protrusions, which extend from the rim like the rings *in vitro* (Fig. 1D). In both instances, the protrusions point toward the kMT plus end, possibly as a result of interactions with other kinetochore proteins. The surface opposite the protrusions is relatively featureless, similar to the rings seen *in vitro*. From these symmetrized rings, we estimate that partial rings missing four or more Dam1C/DASH heterodecamers would have a gap large enough for a MT to pass through them (Fig. 3G).

### Unattached Dam1C/DASH forms partial rings, which remain clustered

Damaged spindles activate the SAC and can cause kinetochore detachment (Gillett et al., 2004). To test how spindle disruption affects kinetochore organization, we treated Dam1p-GFP-expressing cells with the spindle poison nocodazole and then imaged them by immunofluorescence microscopy (Fig. 4A). Excluding a small subset of cells that lacked both Dam1C/DASH and MT fluorescence signals, cells had either zero (18%, n = 41), one (63%, n = 143), or two punctate tubulin signals (13%, n = 30).

**Figure 4.**
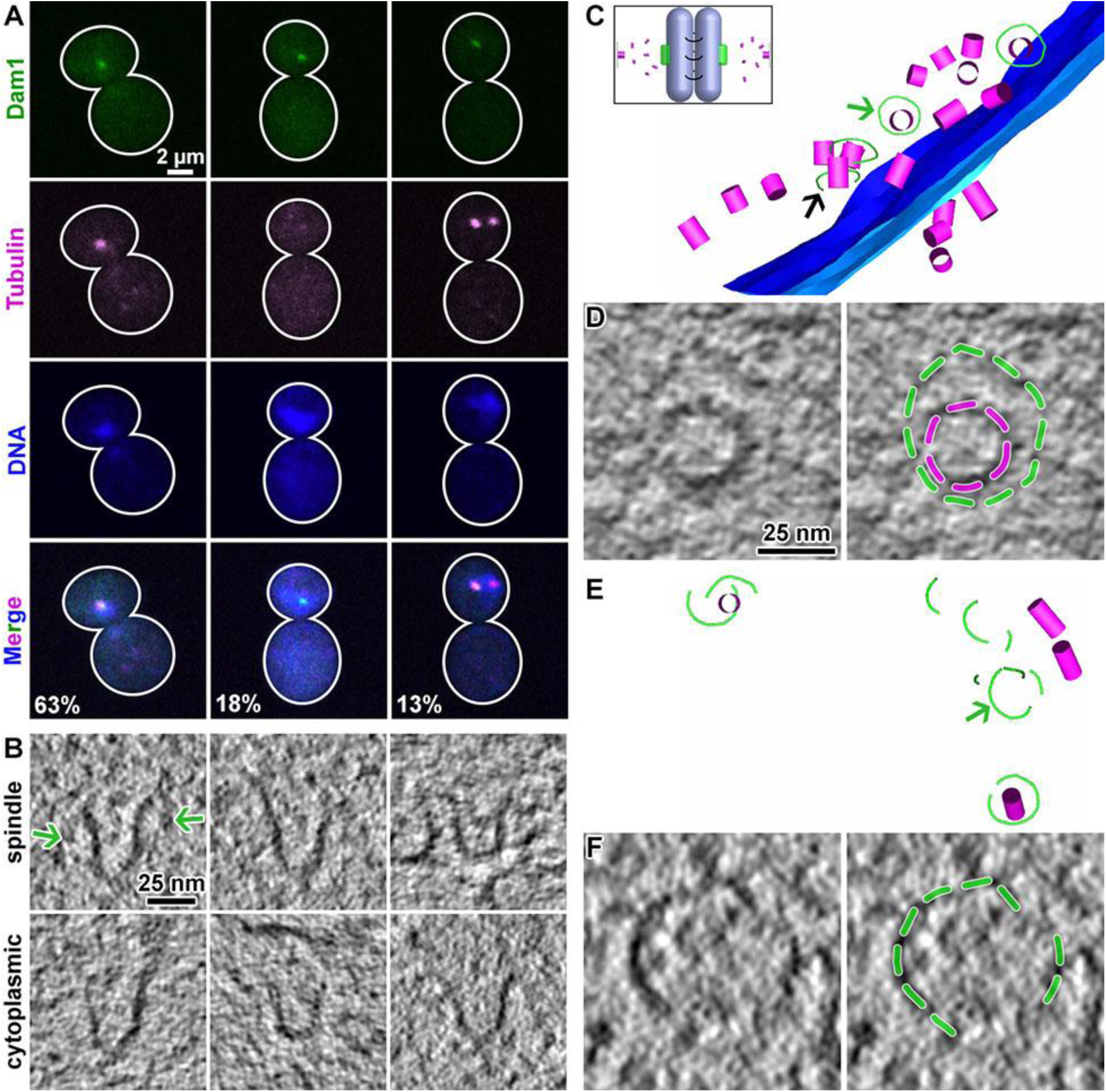
Architecture of spindles in nocodazole-treated cells. Immunofluorescence images of three major spindle morphologies in nocodazole-treated cells. A small minority of the cells had ambiguous morphologies and were not classified. The percentage belonging to each class is printed in the lower row (n = 227). Cryotomographic slices (10 nm) of spindle and cytoplasmic MTs. The plus end is oriented upward in each panel. Green arrows: Dam1C/DASH densities. (C) Model of a nocodazole-disrupted spindle with complete Dam1C/DASH rings associated with short MTs. Part of the bottom-most Dam1C/DASH ring (black arrow) could not be modeled because it was located near the cryosection’s surface. Green arrow: Dam1C/DASH ring that is enlarged in panel D. Inset: schematic showing the effect of nocodazole treatment. (D) Left: cryotomographic slice (8 nm) showing the front view of a complete Dam1C/DASH ring on a short kMT. Right: the same cryotomographic slice but annotated with green dashes over the Dam1C/DASH densities and magenta dashes over the MT densities. (E) Model of unattached Dam1C/DASH oligomers and MT fragments in the nucleoplasm. Green arrow: partial Dam1C/DASH ring that is enlarged in panel F. (F) Left: cryotomographic slice (8 nm) showing unattached Dam1C/DASH partial rings. Right: the same cryotomographic slice but annotated with green dashes overlaying the Dam1C/DASH densities.

Punctate MT fluorescence signals suggests that they are very short and form small clusters. Unlike MTs, all Dam1C/DASH fluorescence was confined to a single focus, suggesting that all the Dam1C/DASH rings formed a single cluster.

The intact spindle is a prominent landmark that facilitated our systematic search for kinetochores in metaphase-arrested cells; this strategy was not possible in nocodazole-treated cells because spindles are disrupted. We therefore performed cryo-ET of 131 randomly chosen cryosections of these cells and were able to locate kMTs in 5 cryotomograms. Consistent with the immunofluorescence data, the nocodazole-treated cells contained small clusters of extremely short MTs (20 - 50 nm long, Fig. 4B and Table S1), all of which had flared ends. In untreated cells, cytoplasmic MTs have their plus ends near the cell membrane, making them extremely challenging to find in cryosections. Owing to their shortness in nocodazole-treated cells, the plus ends of four cytoplasmic MTs were seen in the vicinity of the nuclear envelope. Dam1C/DASH rings were found around some kMTs (nuclear) while cytoplasmic MTs did not have any Dam1C/DASH rings encircling them (Fig. 4B-D). In one instance, we observed the Dam1C/DASH rim in contact with the kMT surface (Fig. 4D), but we did not see any Dam1C/DASH rings contacting the back of the protofilaments’ curved tips. Like in metaphase cells, Dam1C/DASH rings attached to kMT walls via flexible bridges. We also located small clusters of unattached Dam1C/DASH partial rings in the nucleoplasm, far from the spindle pole body (Fig. 4E and F). Our observations are consistent with the notions that kinetochores are clustered by an MT-independent mechanism (Goshima and Yanagida, 2000; Jin et al., 2000; Richmond et al., 2013) and that all sixteen budding-yeast kinetochores work together like a single, much-larger mammalian kinetochore (Aravamudhan et al., 2014; Joglekar et al., 2009; Joglekar et al., 2008). In summary, some Dam1C/DASH subcomplexes detach from damaged spindles and are found as clusters of partial rings. Another subset of Dam1C/DASH rings encircle the extremely short kMTs and only contact the kMT’s flat surface.

### Kinetochore position on the kMT is sensitive to tension

Even if kinetochores are attached to kMTs, the spindle checkpoint can still be activated if tension across the spindle is lost. To determine how the outer kinetochore responds to loss of tension in the presence of attached kinetochores, we imaged metaphase cells in which cohesin can be conditionally cleaved. In these cells, Scc1 is replaced by Scc1-TEV268, which can be cleaved at an internal recognition site by inducible TEV protease (Mirchenko and Uhlmann, 2010; Uhlmann et al., 2000) (and this paper). Light micrographs showed that these cells have a large bud, extra-long spindle, and a multi-lobed nucleus (Fig. 5A). Our cryotomograms confirmed that these cells had distorted nuclei and showed that MTs were absent from the center of the spindle (Fig. 5B and S8). We located 33 Dam1C/DASH rings, which were much more difficult to find because longer kMTs made the plus ends rarer in our cryotomograms and because many rings were located far from the kMT plus end (example serial reconstruction in Fig. 5C and Table S1). Unlike in the other cells we imaged, Dam1C/DASH rings were rarely clustered; only one such cluster was found in this dataset (Fig. S8). The ratio of complete to partial rings in these cells was similar to that in metaphase cells (Table S1).

**Figure 5.**
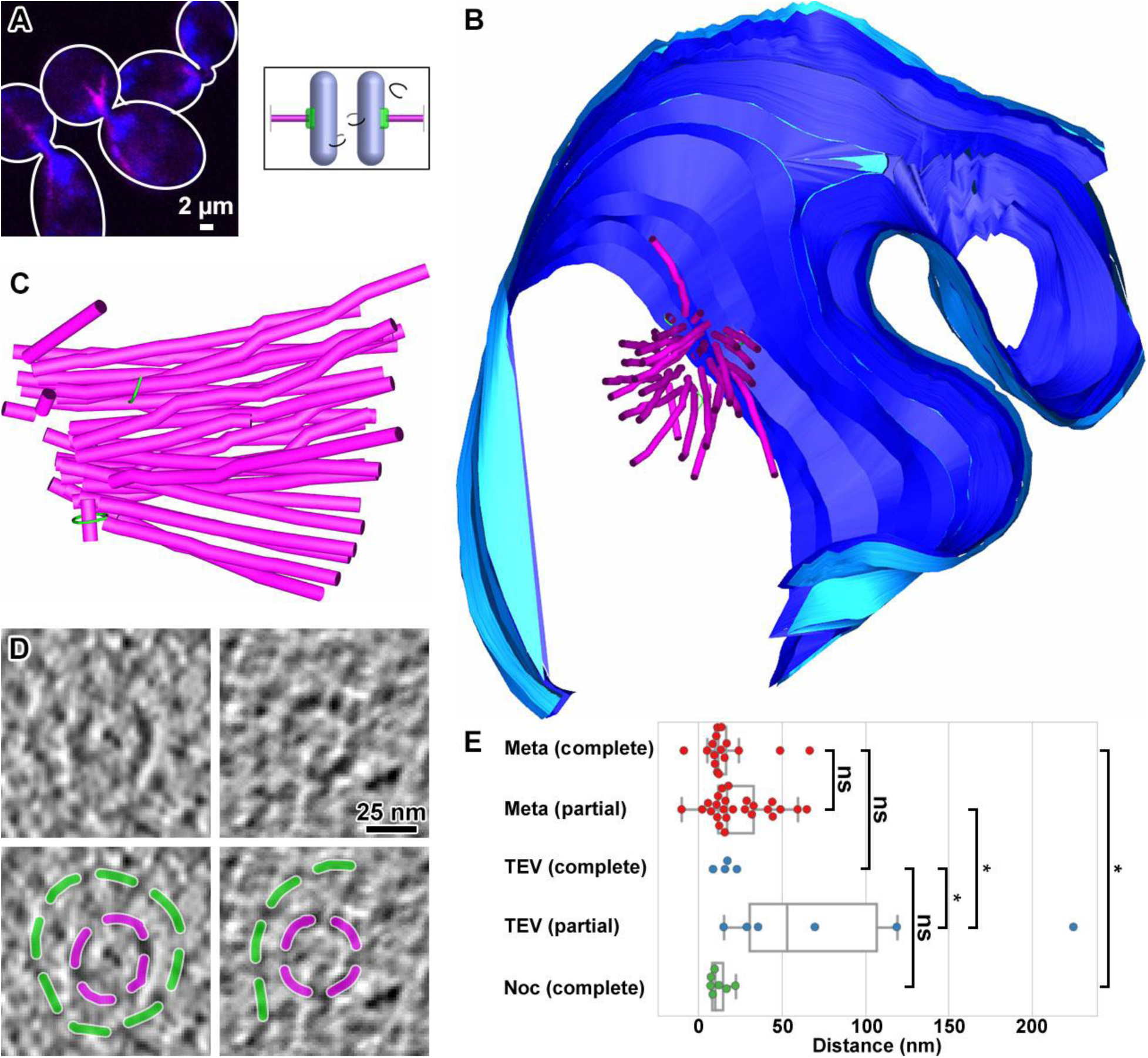
Architecture of spindle machinery in mitotic cells without cohesion. (A) Immunofluorescence image of a metaphase-arrested cell (Cdc20 depleted) in which tension is absent because the cohesin subunit Scc1 is cleaved by TEV protease. Blue: DNA. Magenta: MTs. Inset: schematic showing the loss of cohesion. (B) Serial cryo-ET model of one such cell. The nuclear envelope is colored blue and the spindle MTs colored magenta. The few Dam1C/DASH rings that were found are colored green. Note that the discontinuities in the nuclear envelope model are from the interfaces between adjacent cryosections, which could not be accurately modeled. (C) Enlargement of the spindle modeled in panel B, rotated to a view perpendicular to the spindle’s axis. (D) Cryotomographic slices (6 nm) of front views of complete (left) and partial (right) Dam1C/DASH rings around kMTs. (E) Box-and-whisker plots and raw values (colored circles) of the distances between kMT plus ends and Dam1C/DASH ring centers of mass. Two Dam1C/DASH rings were located in front of kMT plus ends, which gave rise to negative distance values. Meta: cells arrested in metaphase. TEV: cells arrested in metaphase and with Scc1 cleaved. Noc: cells treated with nocodazole. ns: not significant, Student’s t-test p > 0.05. Asterisk: F-test of equal variance p < 0.01.

In the absence of tension, some Dam1C/DASH rings were located very far (> 100 nm) from the kMT plus ends. To test for a correlation between tension and the location of a Dam1C/DASH ring along a kMT, we measured the kMT-tip-to-Dam1C/DASH distance for all three spindle conditions, with complete and partial rings kept as separate groups (Fig. 5E). In metaphase cells with kinetochores under tension, there was no difference between the kMT-tip-to-Dam1C/DASH distances in complete and partial rings (means and standard deviations -- complete ring: 17 ± 18 nm, n = 17; partial ring: 24 ± 18 nm, n = 26; two-tailed t-test p > 0.05). However, in the absence of spindle tension, a few partial rings were located much farther from the kMT plus ends than the complete rings (complete ring: 17 ± 6 nm, n = 4; partial ring: 82 ± 80 nm, n = 6; F-test, p < 0.01). The complete rings in all three spindle states -- metaphase, tensionless, disrupted -- were located close to the kMT plus end (disrupted spindle: 12 ± 6 nm, n = 7, two-tailed t-test p > 0.05 for all pairwise comparisons). In summary, complete rings, but not partial rings, remain associated with the kMT plus end in the absence of tension.

## DISCUSSION

The discovery that the Dam1C/DASH outer-kinetochore complex can form rings around MTs suggested a mechanism for how kinetochores can remain attached to a dynamic kMT tip (Hill, 1985; Miranda et al., 2005). Notably, a kMT-encircling complete ring is thought to be topologically trapped because its means of dissociation is the plus end, which is blocked by the protofilaments’ curved tips. Structural studies have revealed a more complicated picture: Dam1C/DASH can also form spirals and partial rings (Gonen et al., 2012; Wang et al., 2007). Furthermore, calibrated fluorescence-microscopy experiments revealed that each kinetochore has, on average, twelve Dam1C/DASH heterodecamers (Dhatchinamoorthy et al., 2017), challenging the notion that the complete ring is the only functional form of Dam1C/DASH *in vivo*. Our study has now shown that partial and complete partial Dam1C/DASH rings coexist *in vivo*, with partial rings being the majority species. Many partial rings have gaps larger than 25 nm, meaning that those kinetochores do not attach to kMTs by topological means. How would Dam1C/DASH keep chromosomes attached to spindles under tension? We believe that the bridge-kMT interactions *in vivo* are more stable than previously appreciated. In support of this notion, single-molecule studies suggested that Dam1C/DASH oligomers with only one to four heterodecamers, which are not topologically trapped on a MT, can be pulled by depolymerizing MT plus ends (Gestaut et al., 2008). Such stable interactions would be consistent with the observation that the MT-bound Dam1C/DASH pool does not exchange freely with the nucleoplasmic pool (Dhatchinamoorthy et al., 2017).

### Dam1C/DASH is sensitive to both tension and attachment

Spindle integrity and tension at the kinetochore are thought to influence kinetochore structure, leading to SAC signaling. We have experimentally damaged the spindle of some cells and eliminated tension at the kinetochore of others. Our resultant analysis reveals that outer-kinetochore is sensitive to both attachment and tension in these mitotically arrested cells. Dam1C/DASH’s oligomerization state *in vivo* depends on attachment but not tension (Fig. 6A and Table S1). The kinetochore’s position along the kMT’s length is more complicated: they are located near the plus end unless there is no tension and the Dam1C/DASH ring is a partial one. How might these oligomerization and positioning differences be related to the SAC? An early fluorescence-microscopy study showed that in nocodazole-treated cells, kinetochores far from the spindle pole body, but not those nearby, recruited checkpoint proteins (Gillett et al., 2004). Our cryotomograms suggest that in nocodazole-treated cells, checkpoint-protein-associated kinetochores have detached partial Dam1C/DASH rings while the “checkpoint-silent” kinetochores are still attached to short kMTs in the spindle remnant and have complete rings. The causal relationship between the SAC and the Dam1C/DASH phenotypes *in vivo* remain to be determined.

**Figure 6.**
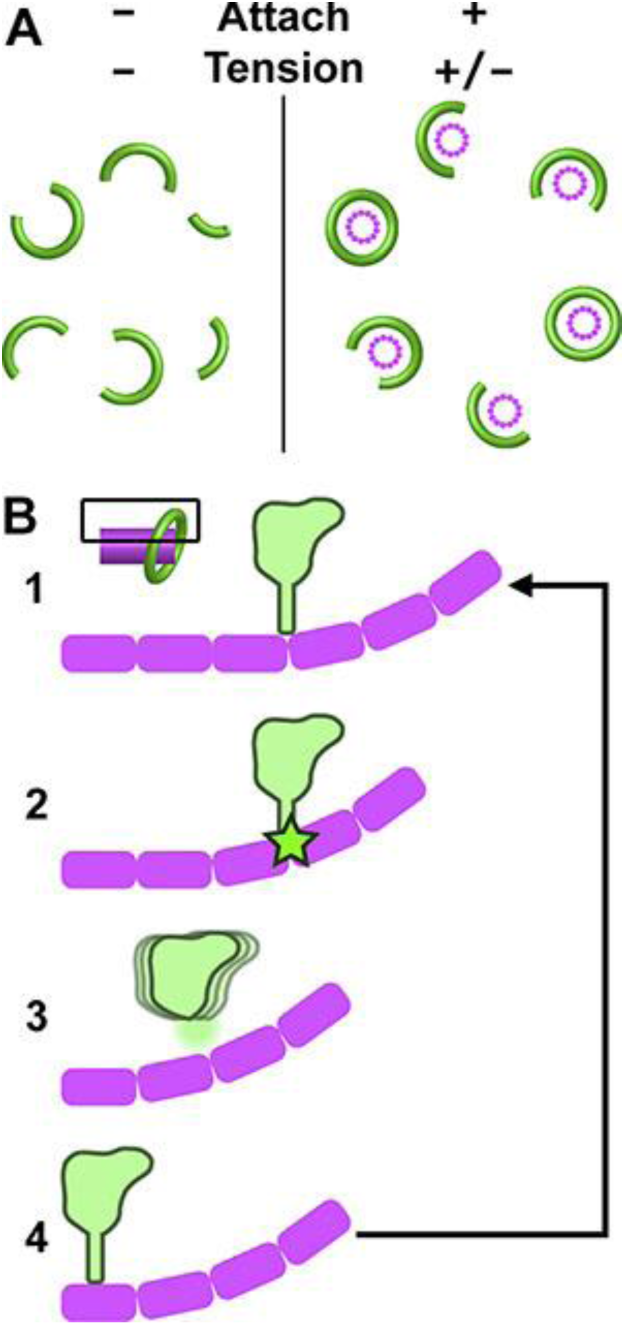
A multi-scale model of the yeast mitotic machinery *in vivo*. Cartoon of clusters of Dam1C/DASH rings, viewed along the spindle axis. Dam1C/DASH (green) can only form complete rings when attached to kMTs (magenta). Inset: cartoon of a single Dam1C/DASH-kinetochore attachment site. The boxed area is enlarged, showing a schematic of Dam1C/DASH in cross section (green) and tubulin dimers (magenta rounded rectangles). (1) The bridge is stably engaged with the flat surface of an MT until (2) the peeling protofilament becomes locally curved enough to destabilize the bridge’s interaction. (3) If enough Dam1C/DASH bridges are freed, the ring can diffuse along the kMT axis until it encounters a flat portion of the MT. (4) Here the bridge makes a stable contact again, attaching Dam1C/DASH to a position closer to the minus end.

### The yeast kinetochore is not a monolithic structure

Dam1C/DASH interacts with the KMN (Knl1, Mtw1, Ndc80) outer-kinetochore network and other kinetochore proteins, many of which have long coiled-coil domains (Caldas and DeLuca, 2014; Wang et al., 2008; Westermann et al., 2005). Such extended structures are skinny and would have been missed in our cryotomograms. However, globular domains such as the Ndc80 calponin-homology domain may account for some of the small densities protruding from kMT plus ends (Fig. 3E). In the vicinity of the kMT, there are also many nucleosome-sized densities, some of which may account for the centromeric nucleosome or the center of the hub-shaped MIND (Mtw1, Nnf1, Nsl1, Dsn1) complex (Dimitrova et al., 2016; Gonen et al., 2012; Tachiwana et al., 2011). Of these complexes, the centromeric nucleosome is expected to be coaxial with the kMT (McIntosh et al., 2013), but we did not observe any enrichment of nucleosome-size densities along this axis. Instead, the majority of the kinetochores probably bind kMTs off-axis *in vivo*, which is a phenotype of purified kinetochores (Gonen et al., 2012). Our cryotomograms are consistent with a model in which the yeast kinetochore is a highly flexible structure and with its mass spread over a large volume (Dimitrova et al., 2016) (Steve Harrison, personal communication].

### A model for microtubule-driven chromosome movement

Yeast chromosomes move poleward along kMTs by two different mechanisms. Newly assembled yeast kinetochores first contact the side of a kMT and slide poleward by means of the kinesin Kar3 (Tanaka et al., 2005). Eventually, the kMT plus end contacts the kinetochore, leading to an “end-on” interaction and kMT-driven chromosome poleward movement (Kitamura et al., 2007; Tanaka et al., 2007; Tanaka et al., 2005). There are two popular models of kMT-driven chromosome poleward movement. In the ratchet model (Hill, 1985), kinetochores attach to the spindle by numerous weak interactions and undergo a random walk along kMTs, but have biased poleward movement by the receding plus end. In the forced-walk model (Efremov et al., 2007), the depolymerizing protofilaments push a strongly bound kinetochore and force it to slide poleward. We did not observe any instance of Dam1C/DASH in contact with the protofilaments’ curved tips in any of the three conditions. Contact between Dam1C/DASH and the protofilaments’ curved tips must therefore be transient. We did frequently observe contacts between the Dam1C/DASH bridge and the MT’s flat surface. To explain these observations, we propose a model that incorporates ideas from both the forced-walk and ratchet models (Fig. 6B). Steps 1 - 2: Once the kMT surface underneath Dam1C/DASH becomes curved enough, bridge detachment is triggered. Step 3: If a sufficient number of Dam1C/DASH heterodecamers lose contact, then the Dam1C/DASH ring can diffuse. Step 4, equivalent to step 1: Once the Dam1C/DASH ring translates to a position where straight protofilaments are available, its bridges can reattach. As the kMT shortens, Dam1C/DASH heterodecamers must cycle between attached and detached states, biased poleward by transient steric interactions between Dam1C/DASH and protofilament curved tips. Human kinetochores may also use this kMT-driven segregation mechanism if the functional homolog of Dam1C/DASH, called Ska1 (Abad et al., 2014; Hanisch et al., 2006; Janczyk et al., 2017; van Hooff et al., 2017; Welburn et al., 2009; Zhang et al., 2017), can switch rapidly between bound and unbound states.

## MATERIALS AND METHODS

### Cell strains

All strains used in this study are detailed in Table S2.

### Cell culture and metaphase arrest

Strain US1375 was grown in 50 ml YEPG (YEP: 10% yeast extract, 20% peptone, supplemented with 2% galactose and 2% raffinose) at 30°C, 250 RPM, to mid-log phase (OD_600_ = 0.5 - 1.0) before a change of growth medium to YEPD (YEP with 2% glucose). All growth-medium changes were done by draining YEPG with a vacuum filter, washing with twice the volume of YEPG, and then resuspending the cells in YEPD. Next, the cells were kept in YEPD at 30°C for 3 hours to arrest at metaphase. Right before self-pressurized freezing, the cells were checked by light microscopy for signs of large buds, which indicates successful metaphase arrest.

### Metaphase arrest without cohesion

Strain US4780 was grown in YEPD without methionine overnight, then arrested in G1 phase by addition of alpha factor to 5 µg/ml. The cells were then washed free of alpha factor and then arrested at metaphase by incubation in YEP + raffinose + methionine medium for 4.5 hours. Metaphase-arrested cells were then incubated in YEPG for 1.5 hours to induce TEV protease expression.

### Nocodazole arrest

Strains US1363 and US8133 were grown overnight in YEPD before arresting at G1 phase by incubation in YEPD containing 0.8 µg/ml alpha factor for 3 hours. The arrested cells were then washed free of alpha factor and released into YEPD containing 15 µg/ml nocodazole. Cells were self-pressurized frozen after 4 hours of incubation.

### Immunofluorescence

Yeast cells were collected by pelleting 1 ml of liquid culture at 15,000 *x g* for 1 minute. The pellet was then fixed in KPF (100 mM K_2_HPO_4_, 4% paraformaldehyde) at 22°C for 1.5 hours. The cells were then washed three times with 1 ml 100 mM K_2_HPO_4_, then once with 1 ml SB (1.2 M sorbitol, 100 mM phosphate-citrate). Next, the cells were incubated at 30°C in 200 µl SB containing glusulase and zymolase for cell wall digestion. The resulting spheroplasts were washed with SB and then incubated with primary antibody (diluted 1:1000) for 2 hours at 22°C. After washing out the unbound primary antibody with BSA-PBS (1% BSA, 40 mM K_2_HPO_4_, 10 mM KH_2_PO_4_, 150 mM NaCl), the spheroplasts were incubated with secondary antibody (diluted 1:2000) for 2 hours at 22°C. After washing out excess secondary antibody with BSA-PBS, the spheroplasts were suspended in 5 µl of mounting media (Vectashield H-1200, Vector Laboratories, Burlingame, CA) and imaged using a Perkin Elmer spinning disc confocal microscope.

### Dam1C/DASH expression, purification and assembly

Dam1C/DASH heterodecamers were expressed and purified using slightly modified published protocols (Miranda et al., 2005; Westermann et al., 2005). All protein buffers contained protease inhibitor (cOmplete, Sigma, St. Louis, MO). The plasmid pC43HSK3H (Miranda et al., 2005) was transformed into BL21 Rosetta 2 (DE3) pLysS cells. This plasmid expresses all ten Dam1C/DASH polypeptides (Dam1p, Dad1p, Dad2p, Dad3p, Dad4p, Duo1p, Ask1p, Spc19p, Spc34p, Hsk3p). Cells were grown to OD_600_ = 0.4, then induced by addition of IPTG to 1 mM. After 4 hours of induction at 37°C, the cells were pelleted by centrifugation at 5,000 *x g* for 15 minutes. The cells were resuspended in 30 ml sonication buffer (20 mM sodium phosphate pH 6.8, 500 mM NaCl, 1 mM EDTA, 20 mM Imidazole, 0.5% v/v Triton X-100) and lysed by sonication at 4°C for 5 minutes (power: 500 W, frequency: 20 kHz; amplitude: 35%, pulse: 0.5 s, elapsed: 0.5 s). The lysates were then centrifuged at 15,000 *x g* for 30 minutes to remove the debris. Ni-NTA agarose beads (5 ml) were exchanged into sonication buffer by twice pelleting at 100 *x g* for 2 minutes, then resuspending in sonication buffer. The Ni-NTA beads were then mixed with the lysates and incubated at 4°C for 2 hours. Next, the Ni-NTA beads were pelleted by centrifugation at 2,000 *x g* for 2 minutes and washed with sonication buffer twice. Elution buffer (20 ml) was added into the Ni-NTA beads and rotated overnight at 4°C at 200 RPM. The eluate was centrifuged at 100 *x g* for 2 minutes. The supernatant was dialyzed to SP low-salt buffer (20 mM sodium phosphate pH 6.8, 150 mM NaCl, 1 mM EDTA) and concentrated to 1 ml. The concentrated eluate was loaded into a 1 ml HiTrap SP sepharose cation-exchange column. The fraction that eluted in 600 mM NaCl was further purified by gel filtration in a Superose 6 column in Superose buffer, which also functioned as the Dam1C/DASH storage buffer (20 mM sodium phosphate, pH 6.8, 500 mM NaCl, 1 mM EDTA). The largest peak was concentrated to 1 ml using a Vivaspin concentrator (2 ml) and then aliquoted to 10 µl per tube. The aliquots were snap frozen with liquid nitrogen and stored at -80°C (Westermann et al., 2005).

### Preparation of control Dam1C/DASH around MTs

Porcine tubulin (5 mg/ml, T240, Cytoskeleton, Denver, CO) was polymerized into MTs and stabilized with Taxol following a published protocol (Westermann et al., 2005), with modifications. The incubation time was extended to 2 hours. The purified Dam1C/DASH heterodecamers (1 mg/ml) were incubated with Taxol-stabilized MTs (5 mg/ml) for 20 minutes at 22°C. For the cryomicrotomy control, purified Dam1C/DASH heterodecamers (2.3 mg/ml) were incubated with Taxol-stabilized MTs (5 mg/ml) for 30 minutes at 22°C. Then, an equal volume of 80% dextran (M_r_ ~ 6,000) was added to the solution and gently mixed before self-pressurized freezing.

### Plunge freezing and self-pressurized freezing of Dam1C/DASH around MTs

Dam1C/DASH-MT complexes (3 µl) were applied on the carbon side of a Quantifoil 2/2 grid (Quantifoil Micro Tools GmbH, Großlöbichau, Germany). Gold colloids (10-nm, BBI solutions, Cardiff, UK) were added as tomographic alignment fiducials. The colloids (20 µl) were first pelleted and the supernatant was removed. Dam1C/DASH-MT complexes (3 µl) were then mixed with the gold pellet and applied to the EM grid. The grid was blotted for 2 seconds with force 2 and then plunged in liquid ethane using a Vitrobot (Thermo, Waltham, MA), set to 100% humidity at 4°C.

### Self-pressurized freezing

Cells and cryomicrotomy control samples were self-pressurized frozen based on a previous protocol (Yakovlev and Downing, 2011). Arrested cells (50 ml) were pelleted at 5,000 *x g*, 4°C for 5 minutes. The pellet was then resuspended in 1 ml of YEPD or YEPG. The cells were re-pelleted at 3,000 *x g* for 1 minute at 22°C and the supernatant was discarded. Next, the cell pellet was resuspended gently in an equal volume of 50% dextran (M_r_ ~ 6,000). The resulting mixture was drawn into a copper tube (0.45 mm outer diameter) using a syringe-style loading tool (Part 733-1; Engineering Office M. Wohlwend, Sennwald, Switzerland). Both ends of the copper tubes were tightly clamped shut before being dropped into liquid ethane.

### EM grid preparation for cryosections

Parallel-bar grids (G150PB-CU, EMS, Hatfield, PA) with continuous carbon film were used for serial cryo-ET. The grids were plasma cleaned at 15 mA for 45 seconds. To coat the grids with gold fiducials, the carbon side of the grids were covered with 4 µl of a solution containing 0.1 mg/ml BSA and 10-nm gold fiducials in water (BBI). The coated grids were air dried overnight then stored in a dry box until use.

### Cryomicrotomy

All cryomicrotomy was done with a UC7 / FC7 cryomicrotome (Leica Microsystems, Vienna, Austria). The frozen copper tubes were trimmed with a diamond trimming knife (Diatome, Hatfield, PA) until amorphous ice was exposed. The sample was then further trimmed to produce a 130 µm x 55 µm x 30 µm (length x width x height) mesa. Next, 100-nm-thick cryosections were cut from the mesa using a 35° diamond knife (Diatome) to produce a cryoribbon, under the control of a micromanipulator (Ladinsky, 2010). The ribbon was picked up using a fiber tool and carefully placed onto an EM grid, parallel to the long bars like in Fig. 1D, and then attached by charging with a Crion device (Leica). A laser window was sometimes used to flatten the cryoribbon on the grid. The grid was then stored in liquid nitrogen until imaged. Owing to sensitivity of serial cryotomography to cell positions being occluded by contaminants, only the grids that had the minimum amount of ice contamination were used. We also devised new cryotools to further minimize ice contamination and facilitate cryomicrotomy (Ng, in preparation).

### Electron cryotomography of *in vitro* Dam1C/DASH + MT

Tilt series of *in vitro* Dam1C/DASH + MT samples were collected using Tomo4 (Thermo). Tilt series of +60° to -60° with an increment of 2° were collected at cumulative dose of 100 - 130 e/Å^2^. For defocus phase-contrast data, the nominal defocus ranged from -10 µm to -14 µm. For Volta phase-contrast data, the nominal defocus was -0.5 µm. Tomographic reconstructions were done using the IMOD program *Etomo* (Kremer et al., 1996; Mastronarde, 1997; Xiong et al., 2009). Sequential cryotomograms were joined using *Etomo*.

### Serial electron cryotomography

Serial cryo-ET data was collected using Tomo4. First, cryosections were imaged at low magnification (2,878 *x*) to locate positions that showed the nucleus. Next, a single high-magnification (15,678 *x*) projection image was recorded at a dose sufficient (1 - 2 e/A^2^) to determine if that cell position had any spindle MTs. Successive positions centered on the same cell were marked out in the sequential cryosections and saved as targets for tilt series collection. Tilt series of +60° to -60° with an increment of 2° were collected at a cumulative dose of 100 - 130 e/Å^2^. For defocus phase-contrast data, the nominal defocus ranged from -10 to -14 µm. For Volta phase-contrast data, the nominal defocus was -0.5 µm. See Tables S3 and S4 for more details. Tomographic reconstructions and CTF compensation were done using the IMOD program *Etomo* (Kremer et al., 1996; Mastronarde, 1997; Xiong et al., 2009). Sequential cryotomograms were joined using *Etomo*.

### Tomogram 3-D analysis

MTs in each cryotomogram were located manually and then classified by morphology: plus ends were either blunt, tapered, or had a ram’s horn configuration; the MT midsections appeared as tubes; the minus ends were conical. All MT plus-ends were scrutinized for kinetochore structures. To determine the diameter of Dam1C/DASH rings, tomograms were oriented to present the *en face* view of Dam1C/DASH before taking the measurement. To determine distances between kMT plus ends and Dam1C/DASH rings, the tips of kMT plus ends and Dam1C/DASH rings were first treated as two circular disks, then the distance between the center of both disks was taken.

### Rotational symmetry analysis

Rotational power spectra were estimated using the python script *ot_rot-ps.py* (https://github.com/anaphaze/ot-tools). This script calls on EMAN2 routines to calculate correlation coefficients between the original image and copies of the image that were rotated in 1° increments (Tang et al., 2007). This correlation function is then subjected to a 1-D Fourier transform, which can then be inspected for the highest degree of symmetry.

### Template matching of reconstituted Dam1C/DASH rings

PEET was used to automatically find candidate positions of all ring-shaped macromolecular complexes in cryotomograms of Dam1C/DASH reconstituted on MTs (Heumann, 2016). First, a sparse series of model points were seeded in the lumens of MTs that were encircled by Dam1C/DASH rings. Extra points were then automatically added with Andrew Noske’s *curve* tool, implemented in the IMOD program *3dmod*. Two types of reference volumes were tested: 1) a lone featureless 50-nm-diameter ring and 2) this same ring encircling a short featureless 25-nm-diameter tube with 4-nm-thick walls. To minimize the effects of densities from the buffer and especially the MTs that protrude beyond the plane of the ring, the subvolumes were masked with a ~ 13-nm-tall, 60-nm-diameter cylinder that completely encloses the Dam1C/DASH ring. To assess the performance of the template-matching runs, the “save individual aligned particles” option was enabled in PEET. At the end of the search, overlapping hits were automatically removed by the PEET *removeDuplicates* routine. To minimize the number of false negatives, the correlation-cutoff was set to 0.

### Subtomogram classification and averaging

Subtomogram analysis was performed using RELION 2.0 and 2.1 with the 2-D and 3-D classification routines (Bharat and Scheres, 2016; Kimanius et al., 2016). The centers of mass of each template-matching hit were imported in RELION. A preliminary round of 2-D classification did not reveal any “junk” classes, e.g., ice crystals, contaminants, and partial rings, probably because the reference model (ring around a short tube) does not resemble the junk classes found in typical cryo-EM samples (Bharat et al., 2015). All template-matching hits were then subjected to 3-D classification, using a featureless 50-nm-diameter ring as an initial reference. The influence of buffer, MT, and nearby Dam1C/DASH densities was minimized by the application of a “soft” edged lifesaver-shaped mask (15-nm thick, with 18- and 30-nm inner and outer radii, respectively). An initial round of asymmetric 3-D classification revealed complete rings highly tilted to various degrees around the MT, partial rings, and spirals; the latter two classes of Dam1C/DASH assemblies were excluded from subsequent analysis. The remaining classes were very similar and had signs of 17-fold rotational symmetry. Dam1C/DASH rings belonging to the class with the clearest 17-fold symmetry were subjected to 3-D autorefinement, using the same mask as before, and with 17-fold symmetry imposed.

For *in vivo* subtomogram averaging, the two most complete rings with the strongest 17-fold rotational power were aligned to a featureless 50-nm-diameter ring using PEET. Seventeen-fold symmetry was then enforced with the Bsoft program *bsym* (Heymann and Belnap, 2007). A 12-nm thick ring-shaped mask was applied to eliminate the MT and nearby nucleoplasmic densities.

### Figures

All cryotomographic slices were generated with the 3dmod *slicer* tool. Isosurface images were rendered with UCSF Chimera (Pettersen et al., 2004). Cartoons and figure layouts were composed with Adobe Illustrator and Photoshop (Adobe Systems, San Jose, CA).

### Data sharing

The 17-fold-symmetrized subtomogram average of reconstituted Dam1C/DASH rings from Fig. S3A was deposited in the EMDataBank as EMD-6912. The serial cryotomogram that comprise the metaphase spindle from Fig. 3B were deposited in the EMDataBank as EMD-6914. The tilt series for all cryotomograms used to make figures in this paper were deposited in the Electron Microscopy Public Image Archive as EMPIAR-10159.

## ACKNOWLEDGEMENTS

We thank the CBIS microscopy staff for support and training, Gemma An for suggesting the use of parallel-bar grids, Shujun Cai for helping with cryo-EM, Simon Jenni and Steve Harrison for Dam1C/DASH plasmids and sharing results prior to publication, Jeff Yong for advice on chromatography, the Jensen lab for computer access, and members of the Gan group, Jack Johnson, Steve Harrison and Paul Matsudaira for feedback.

CTN, CC, LD, and LG were funded by NUS startups R-154-000-515-133, R-154-000-524-651, and D-E12-303-154-217, an MOE T2 R-154-000-624-112, with equipment support from NUS YIA R-154-000-558-133. HHL and US were funded by the Biomedical Research Council of A*STAR (Agency of Science Technology and Research), Singapore.

## Contributions

CTN - experiments, project design, writing, LD - experiments, CC - project design, experiments, HHL - experiments, JS - training, US - project design, writing, LG - experiments, project design, writing.

## Supplemental Information

**Figure S1.**
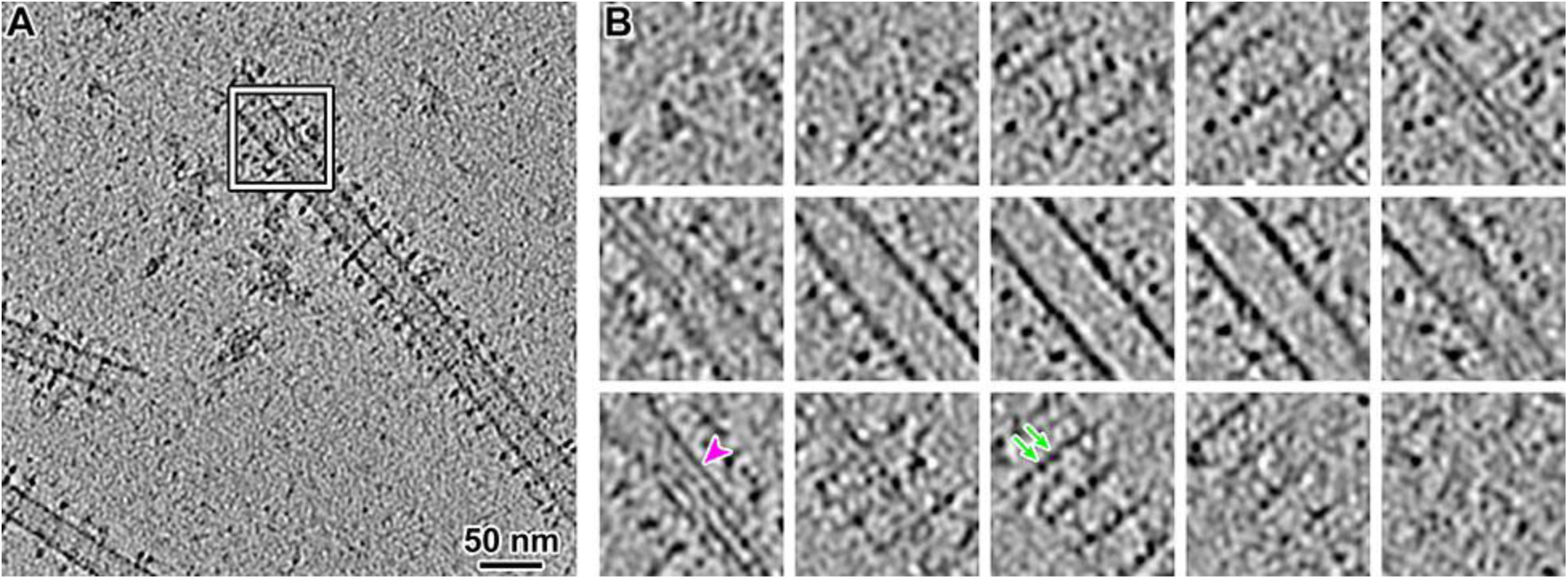
Example plunge-frozen MTs with Dam1C/DASH rings. (A) Cryotomographic slice (60 nm) of MTs, encircled by Dam1C/DASH rings. The amorphous densities below and to the left of the white box are protein aggregates. (B) A series of cryotomographic slices (5 nm) through the position boxed in white in panel A, enlarged twofold. The magenta arrowhead and green arrows indicate a MT protofilament and Dam1C/DASH decamers, respectively.

**Figure S2.**
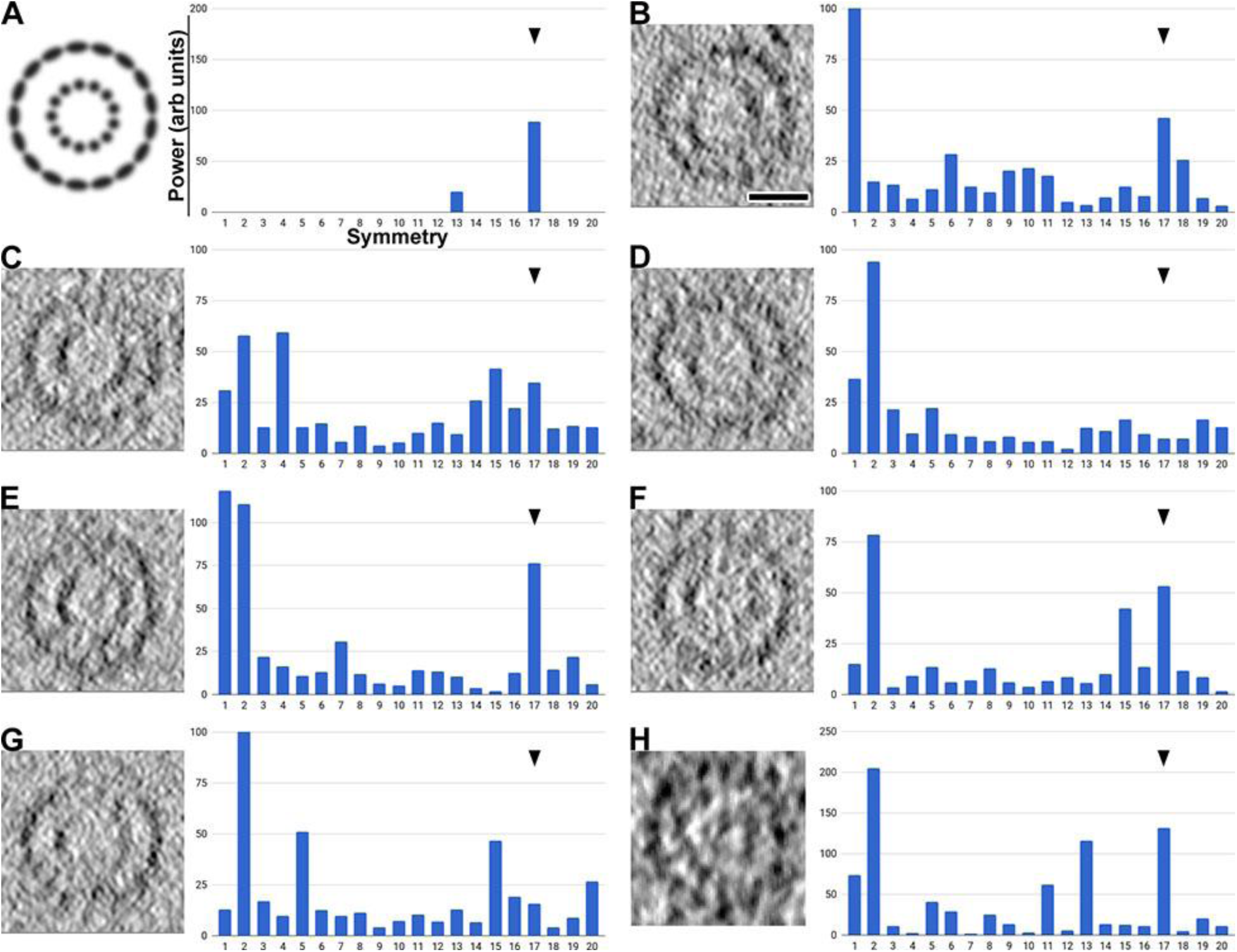
Rotational symmetry analysis of Dam1C/DASH rings *in vivo* and *in vitro*. (A) Left: Manually constructed image with perfect 17-fold (outer) and 13-fold (inner) symmetries. The radii of the outer and inner arrays and their aspect ratios are to the approximate scales of a front view of Dam1C/DASH around MTs. Right: Rotational power spectrum of the densities on the left. The Y axis is the power (arbitrary units) and the X axis is the rotational symmetry. Seventeen-fold symmetry is indicated by the black arrowhead. All subsequent plots have the same axes. (B - G) Left: Cryotomographic slices of Dam1C/DASH ring around MTs *in vitro*, rotated to the front view. Right: Power spectra of the cryotomographic slices. Bar = 25 nm for all cryotomographic slices. The non-Dam1C/DASH densities were masked prior to power spectrum analysis, but the mask is not shown. Some rings, such as that in panels D and G, are distorted and do not produce a strong peak. The 15-fold symmetry peak comes from the MT densities (many MTs have 15-protofilaments *in vitro*), which can leak out of the mask due to the missing-wedge. Note that because these cryotomographic slices were taken coplanar with the Dam1C/DASH ring, the symmetry signal from the MTs are weak or absent when the ring is tilted. (H) Left: cryotomographic slice (6 nm) showing the front view of a Dam1C/DASH ring around a MT *in vivo*. Right: Rotational power spectrum.

**Figure S3.**
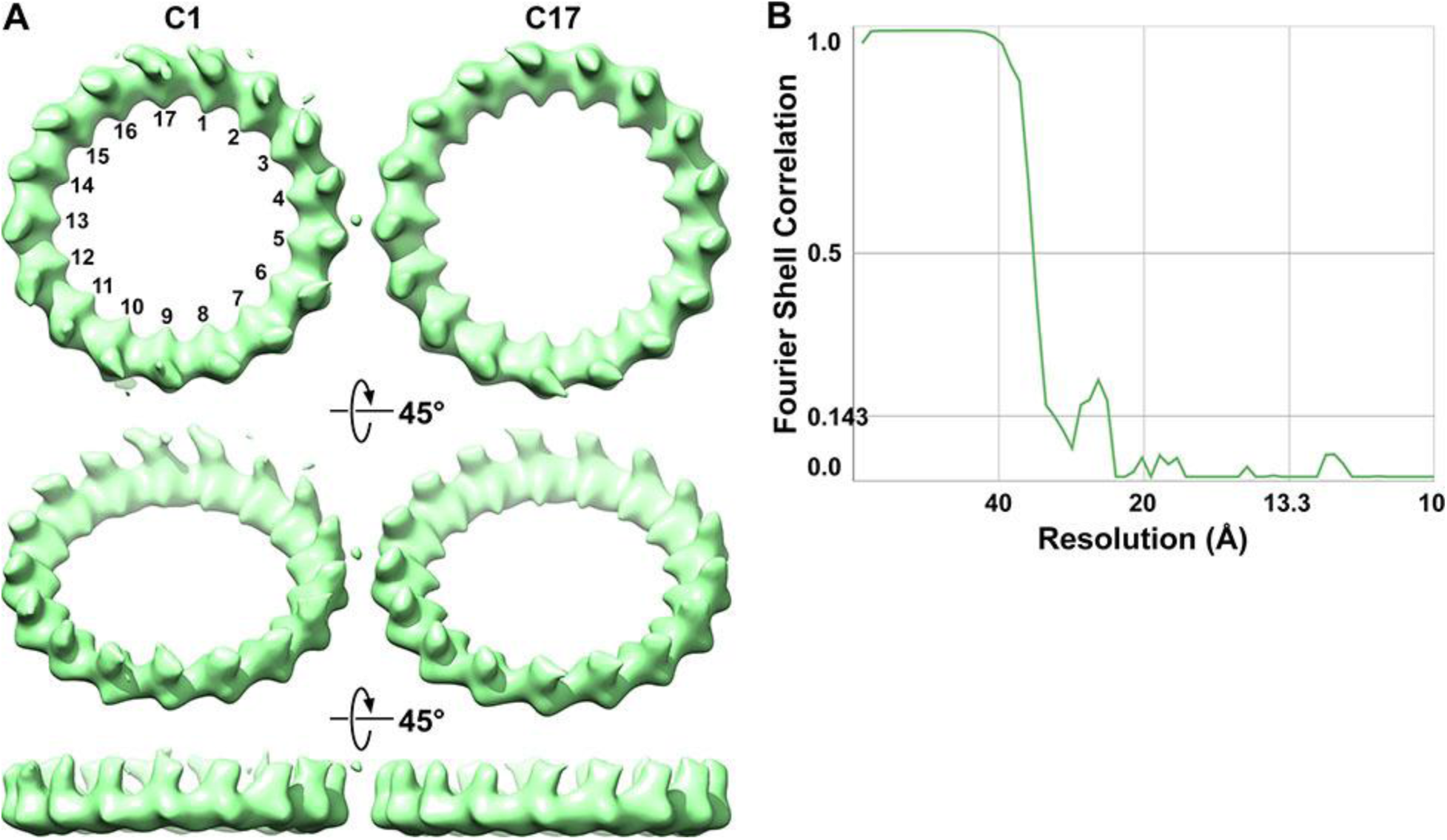
Three-dimensional rotational symmetry analysis of Dam1C/DASH *in vitro*. (A) Subtomogram averages of Dam1C/DASH rings around MTs, without (C1) or with 17-fold (C17) symmetry imposed. The unsymmetrized densities (C1) and subunit numbering are reproduced from Fig. 1D. Only the most symmetric complexes, corresponding to those that resemble Class 1 in Fig. 1, were symmetrized. The upper row is the front view. Each row below is sequentially rotated 45° around the horizontal axis. (B) On the basis of the Fourier-shell correlation = 0.143 criterion, the resolution of the 17-fold symmetrized reconstruction is 32 Å.

**Figure S4.**
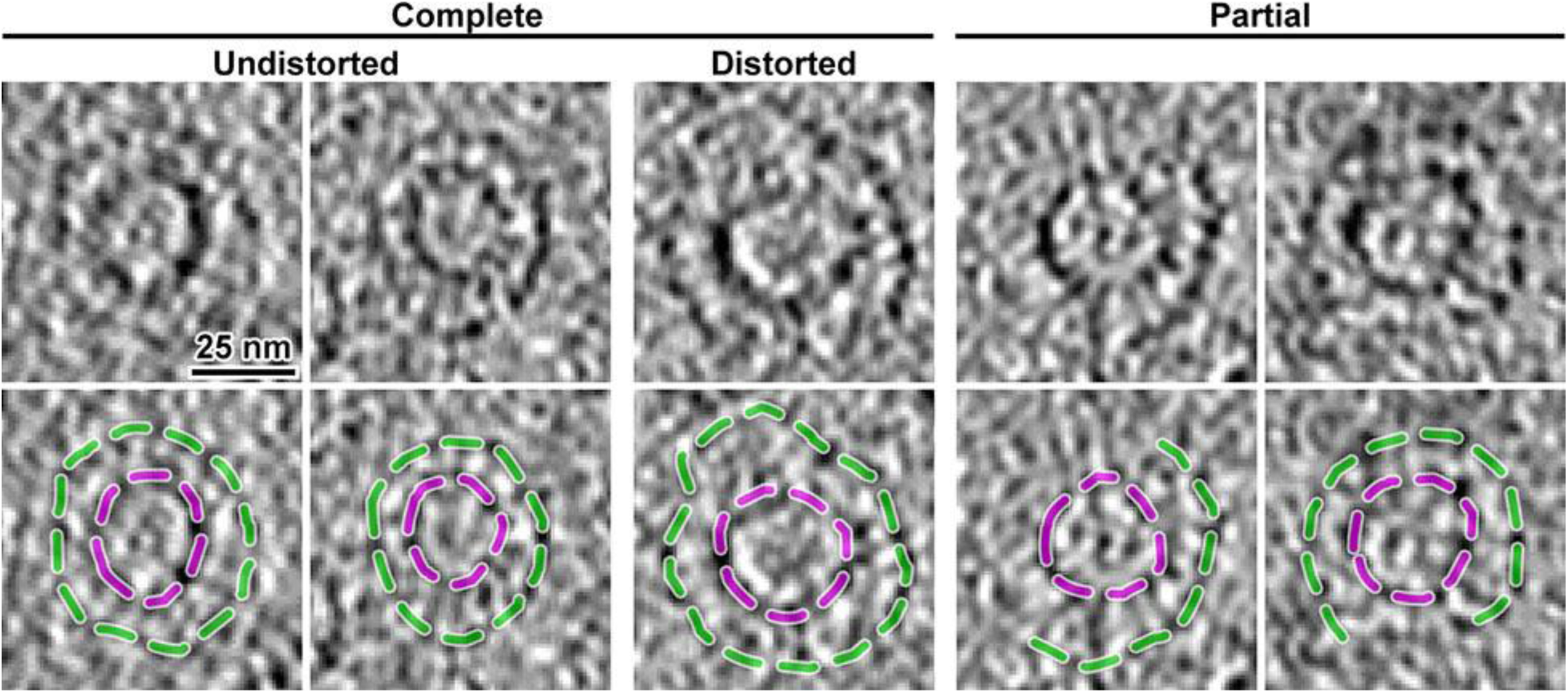
Dam1C/DASH rings can be visualized in cryosections. Dam1C/DASH rings and MTs were assembled *in vitro*, high-pressure frozen, and then cryosectioned. Upper row: cryotomographic slices (6 nm) of Dam1C/DASH rings around MTs. Lower row: dashed lines corresponding to Dam1C/DASH (green) and MT (magenta) densities have been superposed on a copy of the upper panel.

**Figure S5.**
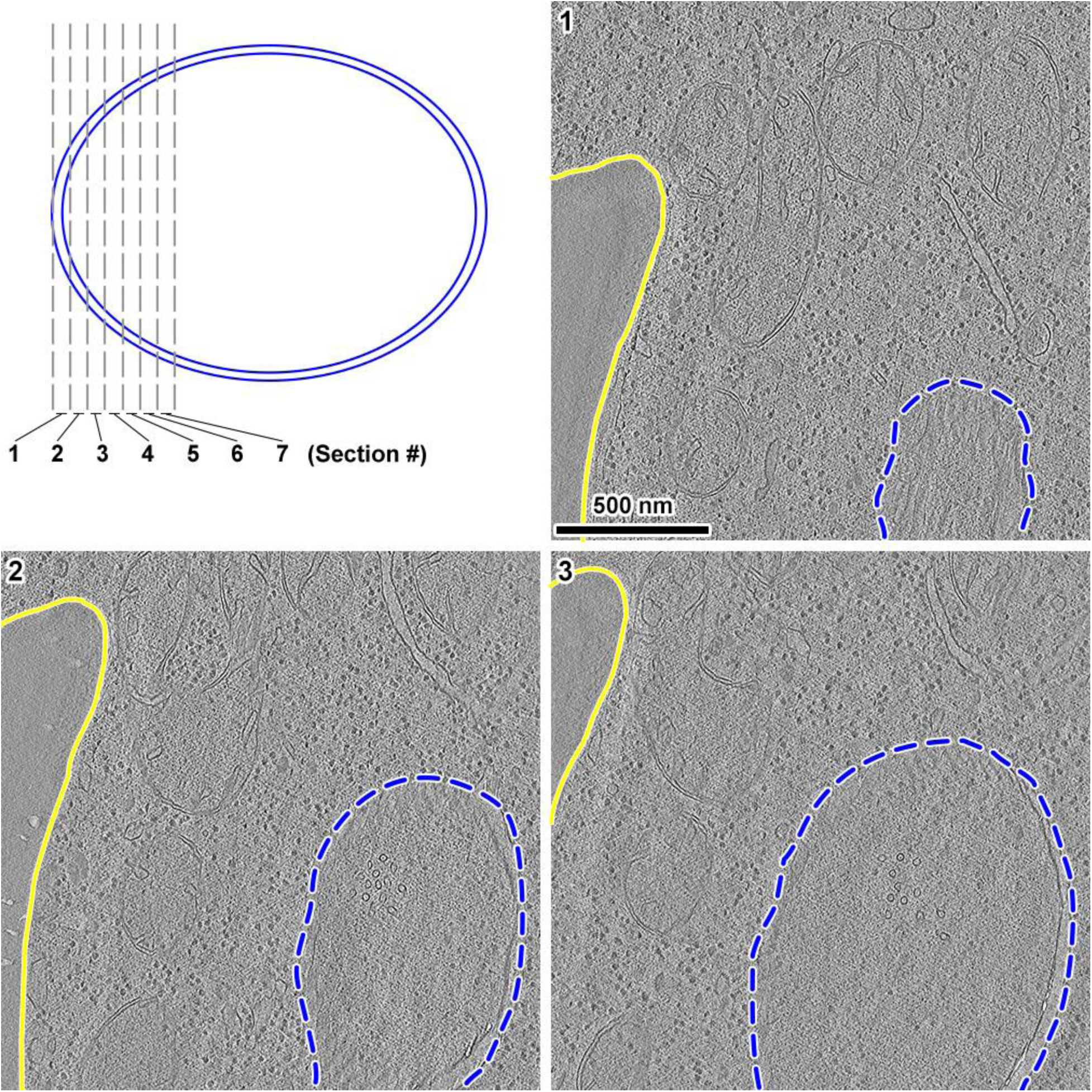
part 1. Serial cryotomograms 1 - 3 of a metaphase yeast cell. (Upper left panel) Cartoon of a cell nucleus, bounded by a nuclear envelope (double blue lines). Seven sequential sections are shown, bordered by vertical gray dashes. Sections are numbered at the upper left of each panel. (Panels 1 - 3) Cryotomographic slices (20 nm) of 3 sequential cryosections of a metaphase cell. The outer nuclear membrane is outlined in blue dashes in each panel. The plasma membrane is outlined by a solid yellow line.

**Figure S5.**
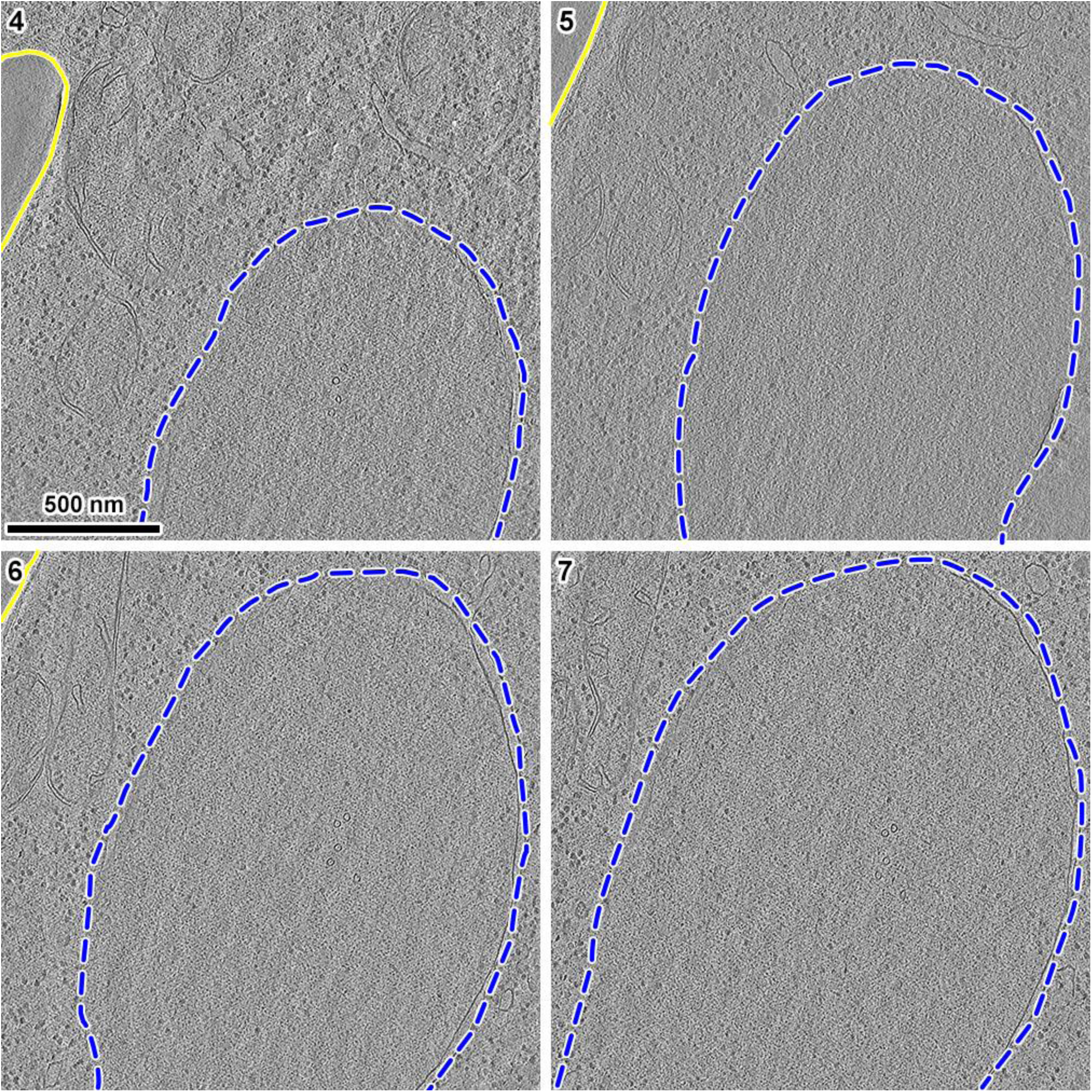
part 2. Serial cryotomograms 4 - 7 of a metaphase yeast cell.

**Figure S6.**
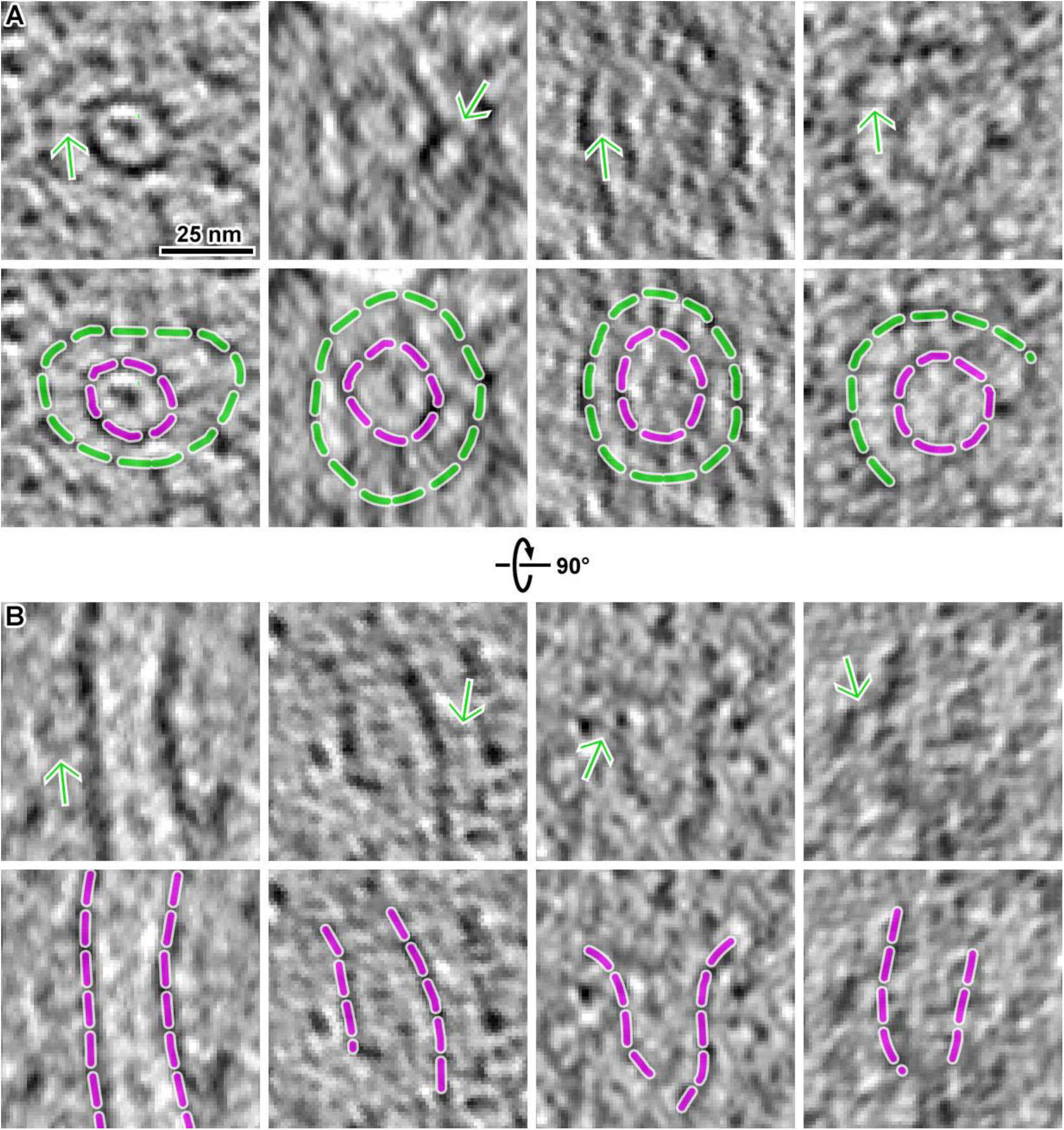
Dam1C/DASH bridges are conformationally heterogeneous *in vivo*. (A) Cryotomographic slices (5 nm) showing four examples of bridges (green arrows) on both complete and partial Dam1C/DASH rings (green dashes) attached to kMT walls (magenta dashes) in metaphase cells. For clarity, the upper and lower panels show the same densities but with different sets of annotations. (B) Same structures as in panel A but rotated 90° around the horizontal axis.

**Figure S7.**
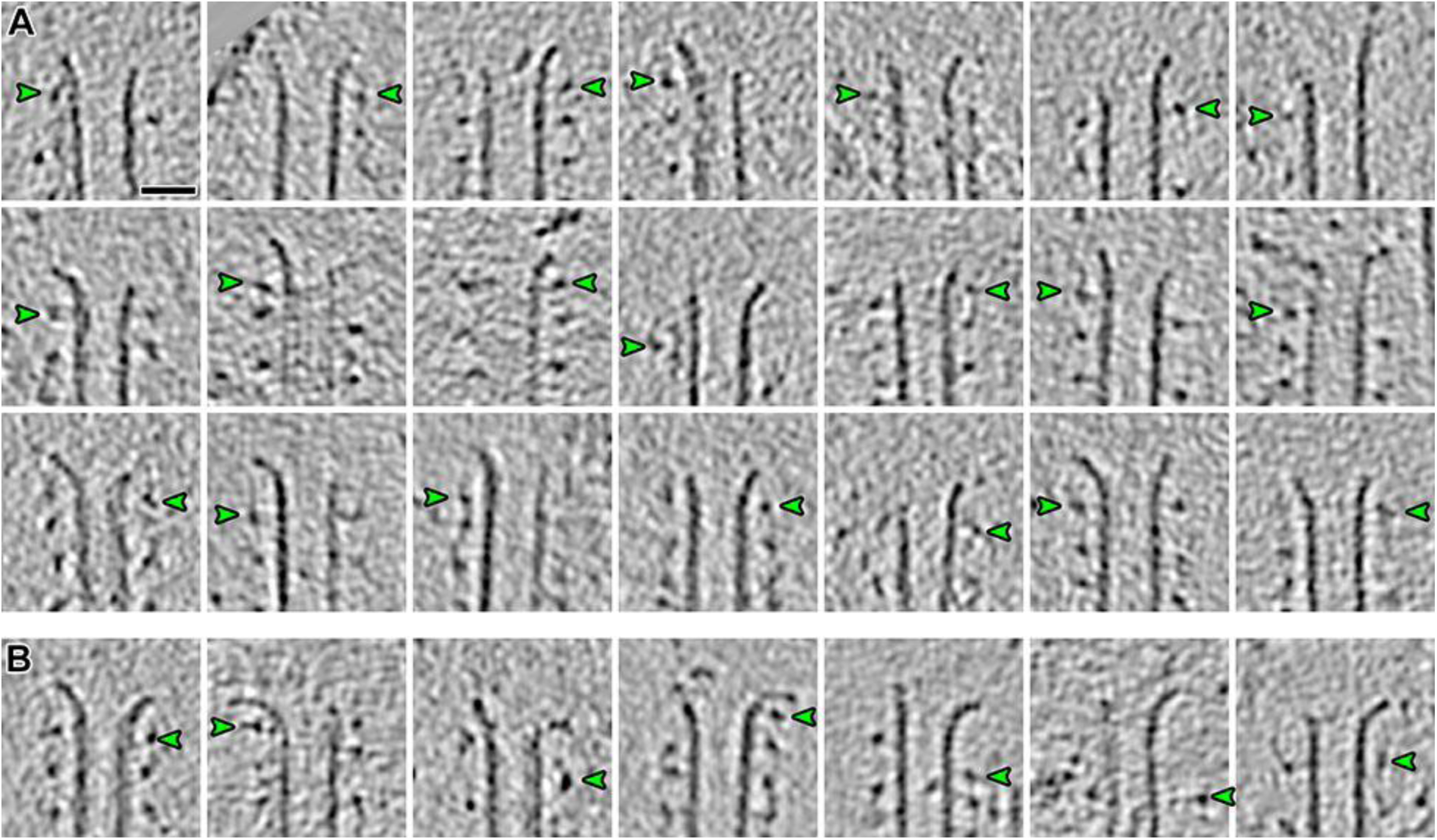
Curved protofilaments rarely contact Dam1C/DASH *in vitro*. (A) Cryotomographic slices (4.6 nm) of the flared ends of MTs assembled with Dam1C/DASH *in vitro*. Green arrowheads indicate the Dam1C/DASH density closest to protofilaments’ curved tip. Scale bar, 25 nm. (B) Same as in panel A but for MTs showing the ram’s horn tip motifs. Note that some MTs appear narrower than 25 nm in a subset of slices taken closer to the surface of the MT. Another subset of MTs have lower contrast because they were oriented almost perpendicular to the tilt axis; this is a well-known missing-wedge effect that changes the appearance of tubular structures.

**Figure S8.**
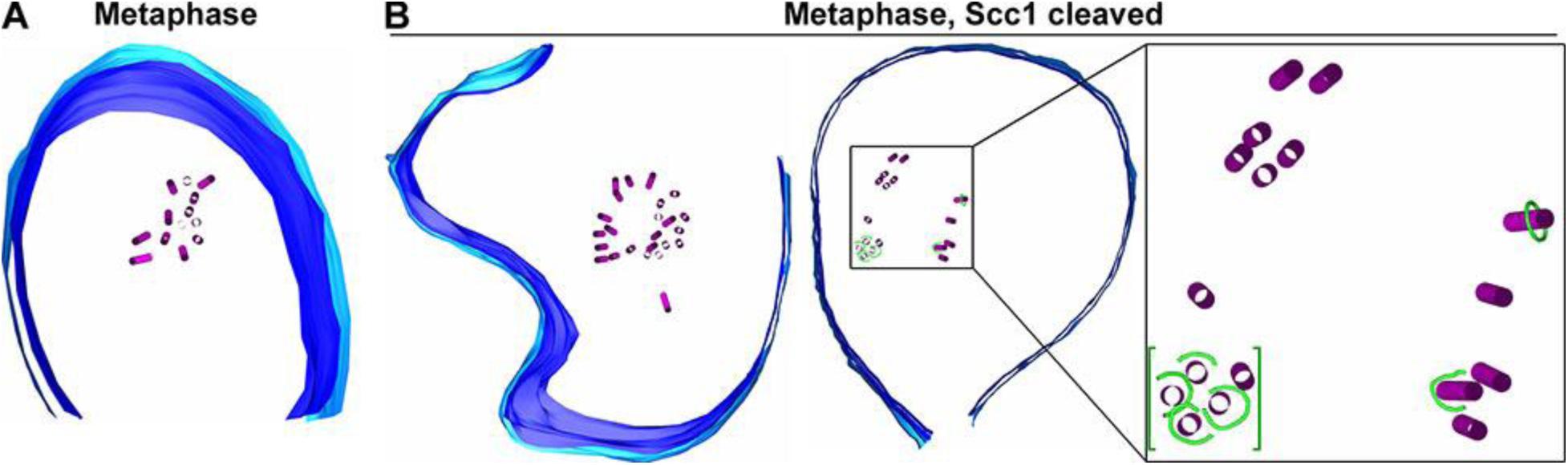
Loss of tension changes the spindle shape and kinetochore distribution. (A) A model of a portion of a metaphase spindle. MTs are magenta and the nuclear envelope membranes are blue. (B) Models of two metaphase spindles with Scc1 cleaved. These models came from single cryosections that were cut almost exactly transverse to the spindle axis. In metaphase cells without tension (Scc1 cleaved), the spindle MTs are arranged in isolated bundles surrounding a MT-free core. The spindle on the right (boxed) includes Dam1C/DASH rings (green) and is enlarged 3.5-fold on the right inset. Note that Dam1C/DASH rings within the cluster (green brackets) were spread out along spindle axis (coming out of the image) and were not in contact.

**Table S1.**
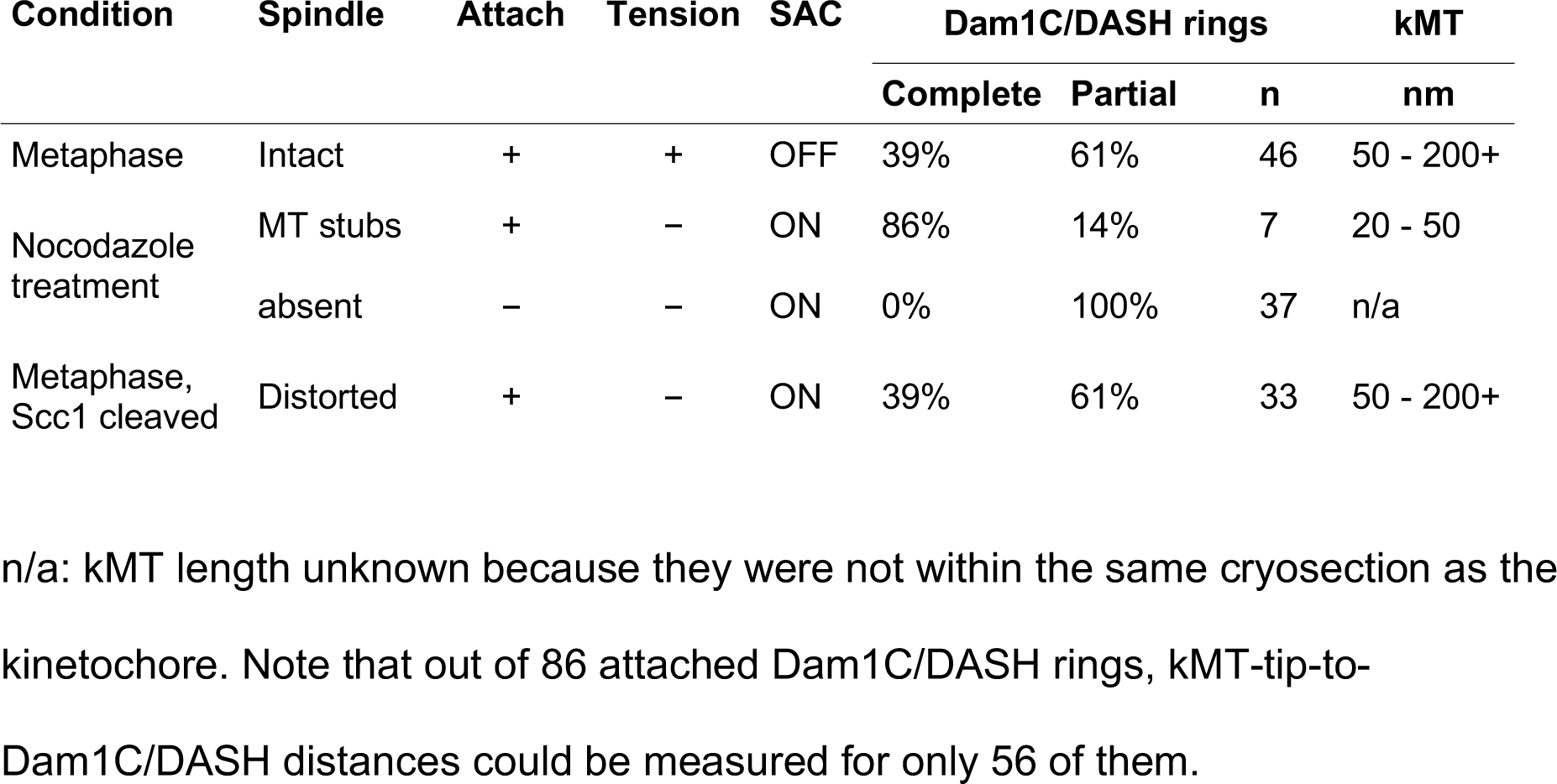
Summary of observations

| Condition | Spindle | Attach | Tension | SAC | Dam1C/DASH rings |  |  | kMT |
| --- | --- | --- | --- | --- | --- | --- | --- | --- |
|  |  |  |  |  | Complete | Partial | n | nm |
| Metaphase | Intact | + | + | OFF | 39% | 61% | 46 | 50 - 200+ |
| Nocodazole treatment | MT stubs | + | - | ON | 86% | 14% | 7 | 20 - 50 |
|  | absent | - | - | ON | 0% | 100% | 37 | n/a |
| Metaphase, Scc1 cleaved | Distorted | + | - | ON | 39% | 61% | 33 | 50 - 200+ |

n/a: kMT length unknown because they were not within the same cryosection as the
kinetochore. Note that out of 86 attached Dam1C/DASH rings, kMT-tip-to-
Dam1C/DASH distances could be measured for only 56 of them.

**Table S2.** Strains used for this study

| Strain | Genotype | Origin | Experiment |
| --- | --- | --- | --- |
| US1375 | MATa ura3 cdc20D: LEU2 his3 GAL-CDC20::TRP1 | Liang, 2012 | Metaphase arrest |
| US1363 | MATa bar1Δ ade2-1 can1-100 leu2-3 his3-11 ura3 trp1-1 (Wild type) | Krishnan, 2004 | Nocodazole treatment |
| US8133 | MATa bar1-1, ura3-1, leu2-3,112, his3-11, can1-100, ade2-1 *Dam1-GFP:TRP1* | This study |  |
| US4780 | MATa, MET3-HA-CDC20: URA3, scc1D: HIS3, SCC1TEV268-HA3-LEU2, GAL-NLS-myc9-TEV protease-NLS2::TRP1, tetR-GFP: HIS3 | This study | Metaphase arrest, Scc1 cleaved |

**Table S3.** Imaging parameters

| Sample | *S. cerevisiae* cells | Dam1C/DASH + MT,<br>cryosections | Dam1C/DASH + MT,<br>plunge frozen |
| --- | --- | --- | --- |
| Grid type | CF-42-2C-T;<br>continuous carbon | CF-22-2C-T<br>(Protochips) | Quantifoil R2/2 |
| Microscope |  | Titan Krios |  |
| Voltage |  | 300 kV |  |
| Gun type |  | FEG |  |
| Camera |  | Falcon II Direct Detector |  |
| Software |  | TOMO4 |  |
| Calibrated<br>magnification | 15,678 / 19,167 | 30,369 | 30,369 |
| Calibrated pixel | 8.93 / 7.3 Å | 4.61 Å | 4.61 Å |
| Defocus | -8 to -15 µm<br>Volta: -0.5 µm | -10 µm | -8 to -14 µm<br>Volta: -0.5 µm |
| Cumulative dose |  | 100 - 130 e/Å<sup>2</sup> |  |
| Dose fractionation |  | 1 / cosine |  |
| Tilt range | ± 60° | ± 60° | ± 66° |
| Tilt increment |  | 2° |  |

## Abbreviations

MT: microtubule
kMT: kinetochore microtubule
cryo-ET: electron cryotomography / cryo-electron tomography

